# Zebrafish differentially process colour across visual space to match natural scenes

**DOI:** 10.1101/230144

**Authors:** Maxime JY Zimmermann, Noora E Nevala, Takeshi Yoshimatsu, Daniel Osorio, Dan-Eric Nilsson, Philipp Berens, Tom Baden

**Affiliations:** School of Life Sciences, University of Sussex, UK; Lund Vision Group, University of Lund, Sweden; Institute of Ophthalmic Research, University of Tübingen, Germany; Bernstein Centre for Computational Neuroscience, University of Tübingen, Germany; Centre for Integrative Neuroscience, University of Tübingen, Germany

## Abstract

Animal eyes evolve to process behaviourally important visual information, but how retinas deal with statistical asymmetries in visual space remains poorly understood. Using hyperspectral imaging in the field, *in-vivo* 2-photon imaging of retinal neurons and anatomy, here we show that larval zebrafish use a highly anisotropic retina to asymmetrically survey their natural visual world. First, different neurons dominate different parts of the eye, and are linked to a systematic shift in inner retinal function: Above the animal, there is little colour in nature and retinal circuits are largely achromatic. Conversely, the lower visual field and horizon are colour-rich, and are predominately surveyed by chromatic and colour-opponent circuits that are spectrally matched to the dominant chromatic axes in nature. Second, above the frontal horizon, a high-gain ultraviolet-system piggy-backs onto retinal circuits, likely to support prey-capture. Our results demonstrate high functional diversity among single genetically and morphologically defined types of neurons.

## Main

Sensory systems have evolved to serve animals’ behavioural requirements. They are tuned to prioritise behaviourally important computations subject to constraints on the neural hardware and metabolic cost (Land & Nilson 2012; Cronin et al. 2014). In vision, specialisations are often made according to the statistics of specific regions in visual space. For example, mouse cones preferentially process dark contrasts above but not below the visual horizon, likely boosting the detection of aerial predators (Baden et al. 2013; Yilmaz & Meister 2013; Calderone & Jacobs 1995). However, beyond anisotropic receptor distributions, systematically linking the statistics of the visual world to the properties of visual systems has been difficult (Lewis & Zhaoping 2006; Ruderman et al. 1998; Webster & Mollon 1997; Buchsbaum & Gottschalk 1983; Simoncelli & Olshausen 2001). Making a link of this kind ideally requires an animal model that allows *in vivo* measurements of light-driven neuronal activity in any part of the eye. In addition, it is necessary to measure the visual characteristics of the animal’s natural world (Dyakova & Nordström 2017), and focus on aspects that are behaviourally important yet sufficiently low-dimensional to be amenable to statistical evaluation. One model that meets these criteria is the colour vision system of the larval zebrafish.

Within three days of hatching, larval zebrafish become highly visual animals with tetrachromatic wide-angle vision (Wong & Dowling 2005; Mehta et al. 2013; Easter, Jr. & Nicola 1996) and well-studied visual behaviours (Preuss et al. 2014; Trivedi & Bollmann 2013; Semmelhack et al. 2014; Temizer et al. 2015; Dunn et al. 2016; Bianco et al. 2011; Muto & Kawakami 2013; Avdesh et al. 2010; Mueller & Neuhauss 2014). Vision is metabolically costly for larval zebrafish: the two eyes make up nearly a quarter of the total body volume, with the neuronal retina taking up >75% of each eye. Indeed, about half of the larva’s central neurons are located inside the eyes (Supplementary Discussion). Space limitations and energy demand create strong evolutionary pressure to make the best use of every visual neuron – potentially driving regional specialisations within the eye. Here, we examine how larval zebrafish retinal circuits process chromatic information in the immediate context of their natural visual world and their behavioural demands. Throughout, we used zebrafish larvae at 7-8 days post fertilisation, *dpf* (for discussion on the choice of age see Supplementary Materials). We find that the eye is functionally and anatomically extremely anisotropic, and these anisotropies match an asymmetrical distribution of colour in the zebrafish natural habitat.

## Chromatic content in nature varies with visual elevation

Zebrafish are surface dwelling freshwater fish of the Indian subcontinent (Arunachalam et al. 2013; Parichy 2015; Spence et al. 2008). Larval and juvenile zebrafish live mostly in shallow, low current pockets on the sides of streams and rice paddies – probably to avoid predation by larger fish (Engeszer et al. 2007), to conserve energy and to facilitate visually guided prey capture of aquatic microorganisms such as paramecia (Semmelhack et al. 2014; Bianco et al. 2011; Preuss et al. 2014; Muto et al. 2017). To systematically record how the visual world varies with elevation in the zebrafish natural habitat we used two complementary approaches: (i) an action camera to take underwater 180° wide-angle, high spatial resolution photographs (Fig. 1-e) and (ii) a custom-built hyperspectral scanner (Baden et al. 2013; Nevala et al. 2017) to take 60° full-spectrum images at a lower spatial resolution, matched to that of the larval zebrafish (Fig. 1f-l). We surveyed n=31 scenes from six field sites in West Bengal, India (SFig. 1a,b, Supplementary Data).

**Figure 1.**
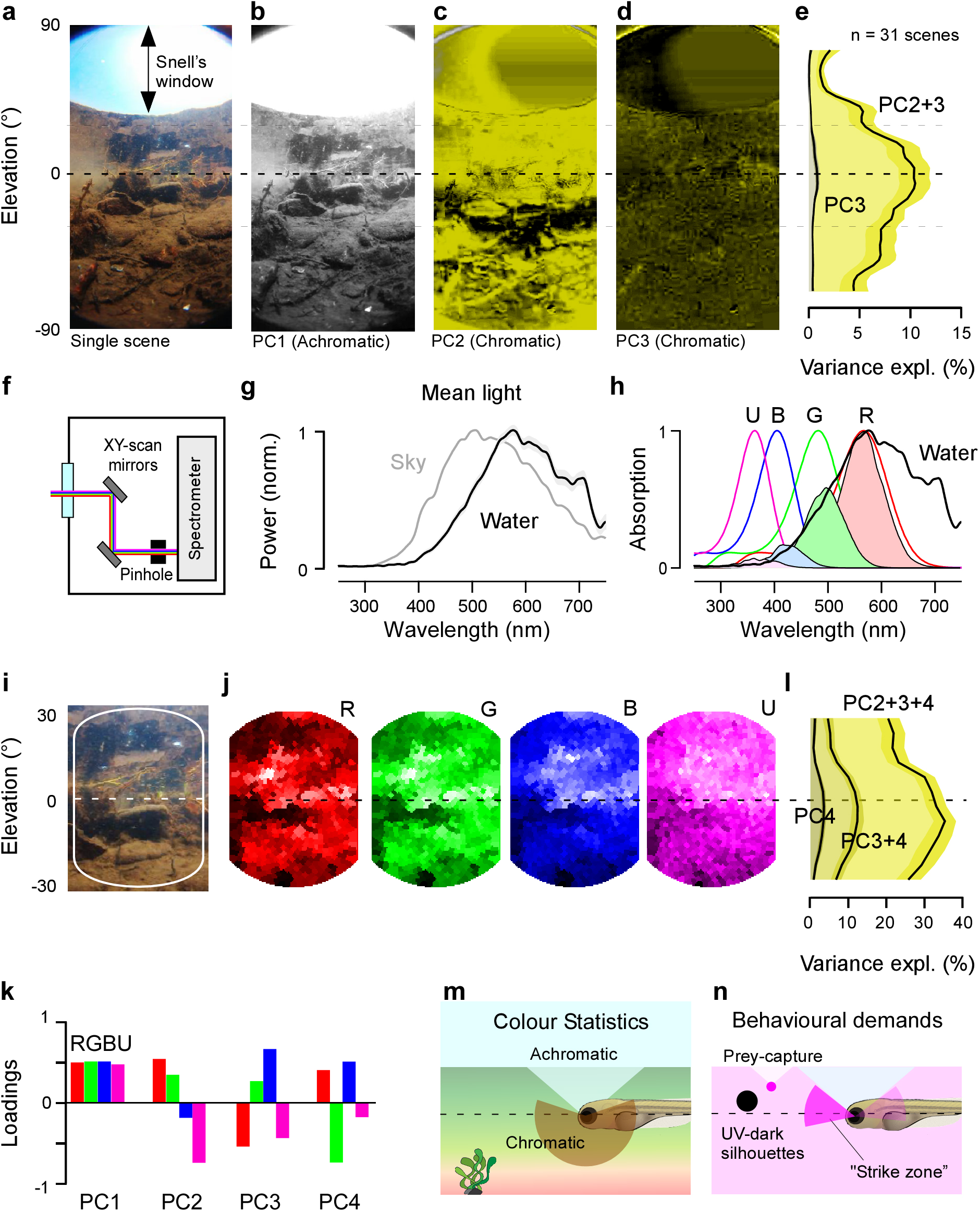
Distribution of chromatic content in the zebrafish natural visual world. **a,** Example 180° underwater photograph taken in zebrafish-inhabited waters in West Bengal, India and **b-d,** the first three principal components across the chromatic dimension (R,G,B) of the image in (a). PC1 reflects the achromatic image content, while PCs 2 and 3 (false-colour coded in shades of yellow) reflect the remaining chromatic content. **e,** Variance explained by PC3 and PC2+3 across all 31 images calculated from 5° horizontal images slices. Error-shadings in s.d.. **f,** Schematic of the custom-built hyperspectral scanner. X and Y mirrors are moved through 1000 regularly spaced positions over a 60° circular window to deflect a ~2.8° spot of light into a spectrometer and thereby build up a hyperspectral image (Baden et al. 2013; Nevala et al. 2017). **g,** Mean of n=31,000 peak-normalised underwater spectra (31 horizon-aligned scenes of 1000 pixels each) and mean spectrum of the sky in zenith above the water. Shading in s.d.. **h,** Zebrafish opsin complement (UV: U, Blue: B, Green: G, red: R), which each opsin-template multiplied with the mean underwater spectrum from (g) to estimate relative photon-catch rates in nature. Templates based on (Baden et al.2013), for discussion on spectral positions see Supplementary Materials. **i,** Enlargement and **j,** reconstruction of photographed scene (a) from scanner-data by multiplying each pixel’s spectrum with each opsin template (h). Scan-reconstructions are truncated beyond 20° from the centre to remove sampling edge artefacts. **k,** Mean loadings of PCs 1-4 across all n=31 scans. **L**, As (e), cumulative variance explained by PCs 2-4 calculated separately for 5° vertical slices across all n=31 scanned scenes. Error-shadings in s.d.. As before (e), most chromatic information exists at and below the horizon. **m,** Schematic summary of natural chromatic statistics and **n,** of expected behaviourally important short-wavelength specific visual requirements. Larval schematic modified from L. Griffiths.

The action camera data demonstrated that in these shallow (<50 cm) waters, both the substrate and the water surface are simultaneously viewed by the larvae’s wide-angle eyes (Fig. 1a), and that the spectrum of light varies strongly with elevation. Directly ahead and to the sides zebrafish eyes align with a mid-wavelength-(“green”) dominated underwater horizon which divides a long-wavelength-(“red”) biased lower visual field and a short-wavelength-(“blue”) biased upper visual field of the ground reflecting the underside of the water surface (Fig. 1a, SFig. 1 b-d). Beyond ~42° elevation, this reflection gives way to Snell’s window (Jerlov 1976; Janssen 1981) – a short-wavelength-biased representation of the 180° world above the water surface compressed into a constant ~97° of visual angle directly above the animal. To estimate which part of the scene contained most chromatic information, we used principal component analysis, using red, green and blue (RGB) pixel values from different elevations. As in terrestrial scenes (Lewis & Zhaoping 2006), PC1 reliably captured the achromatic component where the R, G and B channels co-vary (Fig. 1b). Across the entire image, this component always explained >90% of the total variance.

Next, PC2 and PC3 captured the main chromatic axes (red versus blue; green versus blue) in decreasing order of importance (Fig. 1c,d). Further analysis revealed that the horizon and lower visual field accounted for most chromatic structure, while Snell’s window above the animal was effectively achromatic (Fig. 1e). For this, we horizontally divided each of n=31 images into 5° stripes and calculated the fraction of the total image variance explained by PC2 and PC3 as a function of elevation (Fig. 1e). As our camera was designed for human trichromacy, and can therefore only approximate the spectral content available to the zebrafish’s tetrachromatic retina (Chinen et al. 2003; Endeman et al. 2013), we next computed the chromatic image statistics in hyperspectral images taken at the same sites as seen by the larval zebrafish.

## Spectral positioning of zebrafish cone-opsins under natural light

To sample full-spectrum underwater images in the zebrafish natural world, we custom built a hyperspectral scanner (Baden et al. 2013) comprising a spectrometer and two mirrors mounted on Arduino-controlled servo-motors (Fig. 1f) (Nevala et al. 2017). The system collected 60° full spectrum (200-1000 nm) images centred on the underwater horizon. Individual readings were regularly spaced at ~1.6° to approximate the behavioural resolution limit of larval zebrafish (Haug et al. 2010). A total of 31 scans of 1,000 “pixels” each were taken at the same scenes previously photographed with our action camera (Supplementary Data). To estimate what spectral content is available in nature for zebrafish vision, we multiplied the mean of all 31,000 spectra with the animal’s cone absorption spectra. Zebrafish larvae express mainly four opsins in their four cone types: LWS (548nm), MWS (467 nm), SWS (411 nm) and UVS (365 nm) (Chinen et al. 2003) (for discussion see Supplementary Materials). For simplicity, we will refer to these as the “red” (R), “green” (G), “blue” (B) and “ultraviolet” (U) channels, respectively. As expected (Morris et al. 1995; Chiao et al. 2000), short-wavelengths from the sky illumination were attenuated in the water, resulting in a red-shift of the available light (Fig. 1g). The peak of the mean underwater-spectrum aligned with the absorbance peak of the zebrafish R-opsin (Fig. 1h), suggesting that R-cones are strongly driven in the zebrafish’s natural habitat, and are thus well suited to encode features that require high signal-to-noise representation, such as movement (Schaerer & Neumeyer 1996). In contrast, U-cones lay at the extreme short-wavelength end of available light under water. In this regime, the signal power is ~7% compared to the red channel. Investing neural resources despite the low signal power suggests that zebrafish gain important benefits from using this channel. For example, it could aid detecting UV-rich prey (Novales Flamarique 2012; Flamarique 2016) against the water’s underside internal reflection (Janssen 1981), boost achromatic contrasts against the underwater horizon (Losey et al. 1999; Nava et al. 2011) and more generally to support detection of chromatic contrast. Finally. B-(16%) and G-cones (45%) received intermediate number of photons, and are likely used for both achromatic and chromatic computations alongside the other cones.

## Short-vs-long wavelength computations carry most chromatic information

We next asked which chromatic contrasts in RGBU opsin space predominate in the natural environment of the zebrafish larvae. For this, we multiplied the spectrum of each “pixel” with the spectral sensitivity function of each opsin to yield four monochromatic opsin activation images from each scan (Fig, 1i,j, SFig. 1f, cf. Fig. 1a), one for each opsin channel. As predicted from the available light, the R-opsin image showed the most spatial structure, followed by G and B. In contrast, the U-opsin image had a “foggy” appearance, probably due to UV-light scattering on dissolved organic matter. Such UV-background light can be exploited by animals to detect the UV-dark silhouettes of otherwise difficult to spot objects (Cronin & Bok 2016; Losey et al. 1999).

Next, to separate achromatic and chromatic content across these four opsin images, we computed PCA across now 4-dimensional RGBU opsin space (like above). This again reliably extracted achromatic luminance information into PC1 and then three chromatic dimensions (PC2-4) (Fig. 1k, SFig. 1e). The mean opsin contrasts obtained by PCA across all n=31 scans were (i) RG / BU (“long vs. short wavelength opponency”), (ii) RU / GB and. (iii) RB / GU (complex opponencies). We again cut the images into 5° horizontal stripes and found that the sum of variance explained by the three chromatic dimensions peaked at and below the underwater horizon (Fig. 1l, cf. Fig. 1e).

The efficient coding hypothesis (Attneave 1954; Barlow 1961; Simoncelli & Olshausen 2001) predicts that the obtained opsin contrasts should also be encoded by retinal neurons, as is the case for human trichromacy (Buchsbaum & Gottschalk 1983; Lewis & Zhaoping 2006). Moreover, these circuits should be biased to retinal regions that survey the horizon and lower visual field, where these chromatic contrasts predominate (Fig. 1m). In addition, species specific visual demands that cannot emerge from the statistics of static scenes, such as the need for prey capture and to avoid predators, may drive the evolution of additional, dedicated circuits. For example, zebrafish larvae feed on “translucent” unicellular paramecia that scatter light in a UV-biased manner (Spence et al. 2007; Novales Flamarique 2012; Flamarique 2016). For capture, larvae approach their prey from slightly below and converge the eyes to bring their image into binocular space in front of and just above the horizon (Bianco et al. 2011; Patterson et al. 2013), but outside Snell’s window (J Semmelhack, personal communication). In this part of visual space, the paramecium is illuminated by its own Snell’s window and thus - from point of view of the larval zebrafish - broadcasts its position as a UV-bright spot against the underside of the water (Fig. 1n) (Janssen 1981). For surveying this part of visual space (here dubbed “strike zone”, SZ), the larval retina should invest in UV-driven prey capture circuits. Finally, detecting UV-dark silhouettes against a UV-bright background should work across the entire upper visual field including Snell’s window.

Our natural image data and known behavioural demands of larval zebrafish lead to three predictions for how these animals’ retinal circuits should be organised for efficient coding:

1. Above the animal, light is short-wavelength biased and there is little colour information, but the visual input can be used to spot silhouettes – accordingly, circuits should be achromatic or short-wavelength biased.
2. In the strike zone, the behavioural requirement for prey capture should drive an increased predominance of UV-On circuits.
3. Along the horizon, and below the animal, the retina should invest in chromatic circuits, with an emphasis on short- versus long-wavelength computations. We next set out to test these predictions experimentally. We first assess the distributions of retinal neurons across the eye, and subsequently used *in-vivo* functional imaging to study the chromatic organisation of the inner retina.

## Anisotropic photoreceptor distributions match the distribution of natural light

To study the distribution of the zebrafish larvae’s four cone- and one rod-photoreceptor types across the retinal surface, we fluorescently labelled individual photoreceptor populations. For R-, B- and U-cones we expressed fluorescent proteins under cone-type specific promotors *<u>thrb</u>, sws2* and *sws1*, respectively. No line exclusively labelling G-cones was available. Instead, we calculated their distribution by subtracting genetically defined R-cones from retinae where both R- and G-cones were labelled using immunohistochemistry (zpr-1 antibody (Larison & Bremiller 1990)). Finally, rod-photoreceptors (rods) were surveyed by expressing mCherry under rod specific promoter *xops* (Fadool 2003). We projected each 3D retina as imaged under a confocal microscope into a local-distance-preserving 2D plane, counted photoreceptors and projected their positions back onto the original semi-sphere to generate density maps of each photoreceptor type across the 3D eye (Fig. 2a, Methods).

**Figure 2.**
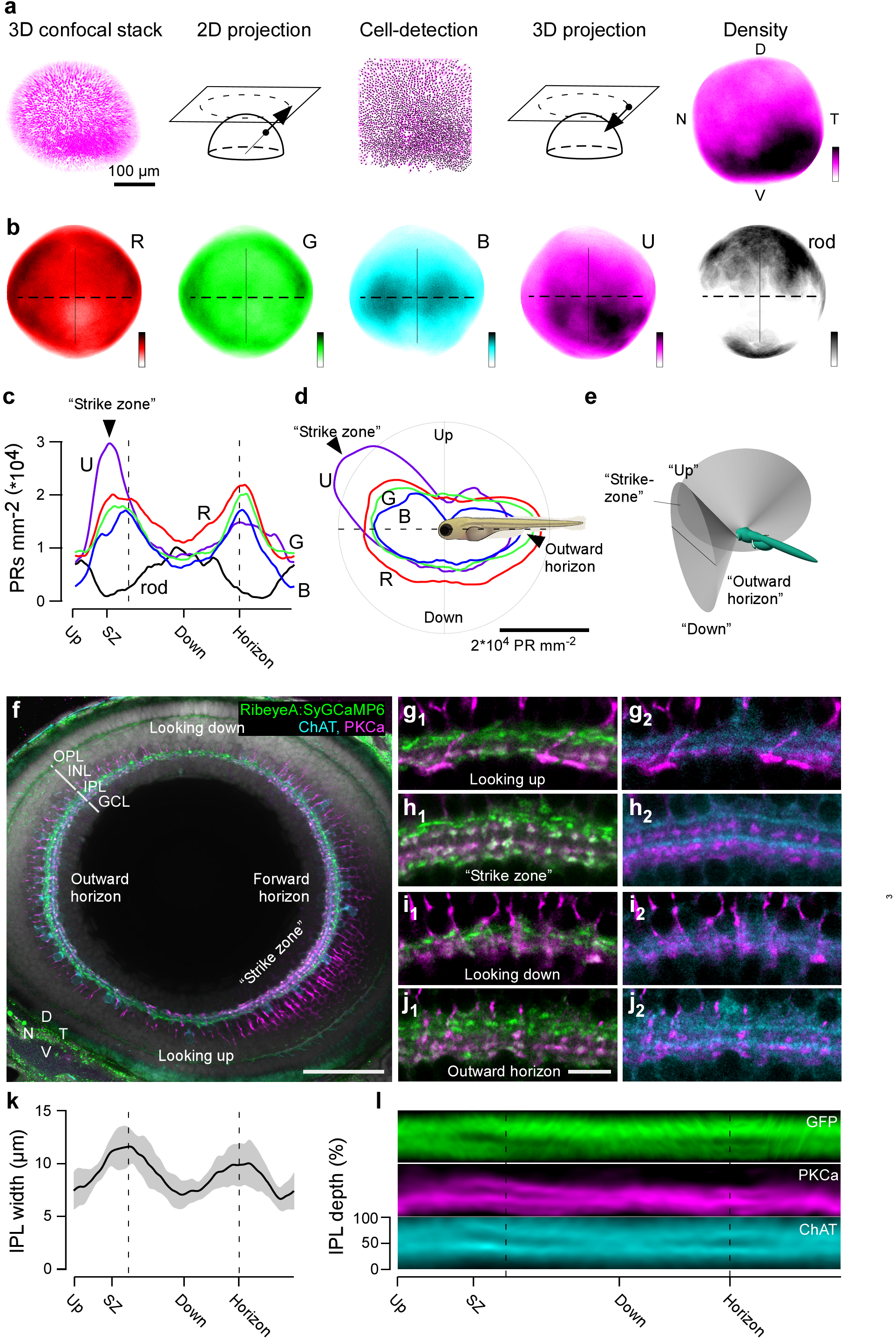
Anisotropic retinal structure. **a,** 3D confocal stack taken across the entire retina of an 8 *dpf* larva *(Tg(Opn1sw1:GFP))* with all UV-cones fluorescently labelled was used for semi-automated quantification of cone densities across the retina (Methods). **b,** Average densities of all four cone types and rods across the retina, based on n=6 (R), 6 (G), 5 (B), 5 (U), 4 (rods) retinas *(Tg(thrb:Tomato)* for R, zpr-1 antibody staining for G, *Tg(-3.5opn1sw2:mCherry)* for B, *Tg(opn1sw1:GFP)* for U, and *Tg(xops:ntr-mCherry)* for rod). Colour Scales: 0 (white) – 35,000 (black) cones mm^-2^ or 0-9,000 rods mm^-2^. D, Dorsal, T, Temporal, V, Ventral. N, Nasal. **c-e**, To compare cone distributions across retinal positions (c,d), we computed densities at sagittal plane approximately aligned with the back-surface of the lens (e, Methods). This plane projects in a ~130° cone from the eye centre, with eyes rotated ~36.5° forward during prey capture (18.5° at rest, SFig. 2a,b). Cone and rod densities across the plane defined in (e) on a linear scale of the fish’s egocentric visual field. Dashed lines indicate the forward and outward horizon. (d) is as (c), plotted in polar coordinates relative to the body of the fish as indicated. **f,** Whole-eye immunostaining of an 8 *dpf Tg(-1.8ctbp2:SyGCaMP6)* larvae labelled against green-fluorescent protein (GFP, bipolar cell (BC) terminals, green), choline acetyltransferase (ChAT, cholinergic amacrine cells, cyan) and protein kinase C alpha (PKCα, On-BCs, magenta). Shown is the same sagittal section used in (f-h). Scalebar 50 μm. OPL, outer plexiform layer; INL, inner nuclear layer; IPL, inner plexiform layer; GCL, ganglion cell layer. **g-j,** Higher magnification sections from (f) showing the IPL at different positions across the eye as indicated. For clarity, GFP/PKCα (g_1_-j_1_) and ChAT/PKCα (g_2_-j_2_) are shown separately. Scalebar 5 μm. **k,** Mean IPL thickness across n=5 whole-eye immunostainings as in (f) and **L,** mean signal in the three fluorescence channels as above.

Unlike in adults, who feature a crystalline photoreceptor mosaic (Engström 1960), in larvae all photoreceptor distributions were anisotropic (Fig. 2b-d). The sum of all cones, which made up ~92% of all photoreceptors, peaked at the horizon (Fig. 2c,d), in line with this part of visual space comprising most chromatic information in nature (Fig. 1e,i). This bias was mainly driven by R-, G-, and B-cones. In contrast, UV-cones peaked ~30° above the forward facing horizon to form a UV-specialised area centralis (Schmitt & Dowling 1999), likely to support visual prey capture (Fig. 1n). Next, the lower visual field was dominated by R-cones, yet closely followed by approximately matched densities all other cones. Like the horizon, this part of visual space could therefore be used for colour vision (Fig. 1m), but with an additional long-wavelength bias as observed in nature (SFig. 1d). In contrast, the upper visual field comprised fewest cones of any type, but instead had an increased number of rods. Unlike cones, rods near exclusively looked straight up through the effectively achromatic but bright Snell’s window, or straight down, perhaps to support the detection of optic flow on the ground even in dim light and/or allow telling the distance to the ground for maximal camouflage for predation from above. Accordingly, already at the level of photoreceptor distributions, the retina of larval zebrafish exhibits a series of anisotropies that align well with the major spectral trends in their visual world and their behavioural demands. How are these anisotropies reflected in the inner retina?

## An anisotropic inner retina

To survey inner retinal structure, we immunolabelled the intact eyes of 7-8 *dpf Tg(-1.8ctbp2:SyGCaMP6)* larvae against GFP (green, all bipolar cell (BC) terminals (Johnston et al. 2014), ChAT (blue, starburst amacrine cells, SACs (Famiglietti 1983; Nevin et al. 2008)) and PKCα (magenta, “On” BCs (Nevin et al. 2008)) and imaged them across the sagittal plane aligned with the back-surface of the lens in the 3D eye (Methods). In larval zebrafish, the full monocular field of view is ~163°, and at rest eyes are rotated forwards by ~18.5° (~35.5° during prey-capture) (Bianco et al. 2011; Patterson et al. 2013). The surveyed sagittal plane samples the visual world in a cone of ~130° (Fig. 2e, SFig. 2), such that its temporal extreme projects in front of the fish while the nasal extreme projects outwards and backwards along the horizon. Dorsal and ventral positions survey the visual world at ~65° elevation below and above the animal, respectively. For simplicity, all visual field coordinates are given in egocentric space from the point of view of the fish: up (ventral), “strike zone” (SZ, temporo-ventral), down (dorsal) and outward horizon (nasal).

Our data on inner retinal structure consolidated and extended all large-scale anisotropies set-up by the photoreceptors (Fig. 2f-l). Like cone-densities (Fig. 2c,d), also inner retinal thickness varied nearly two-fold with position, with the thickest inner plexiform layer (IPL) segments aligning with the horizons (Fig. 2f,k). Alongside, the number, distribution, shapes and sizes of synaptic terminals varied with eye position. For example, while PKCα labelling highlighted three strata in the strike zone (one between and two below the ChAT bands, Fig. 2h_2_), circuits surveying the world above the animal appeared to have only the two lower strata (Fig. 2i_2_). Here, the lowest band featured particularly large BC terminals that are characteristic for teleost “mixed” BCs that process inputs from rod-photoreceptors (Stell 1967) – in agreement with the anisotropic distribution of rods (Fig. 2c). In addition, there were PKCα negative terminals at the IPL bottom that were restricted to the strike zone (Fig. 2h, l). SAC processes also varied with position. For example, the neat bilayer of ChAT immunoreactivity in the strike zone and for looking outward (Figs. 2h_2_, j_2_) disappeared into a “haze” in circuits looking down (Fig. 2i_2_, Fig. 2l). Clearly, the larval zebrafish retina features a broad range of anatomical specialisations across the visual field. How are these anatomical specialisations reflected in function? To address this question, we next turned to calcium imaging of BC terminals across the eye.

## The inner retina is divided into anisotropic chromatic and achromatic layers

We used 2-photon *in vivo* imaging of light-driven activity in retinal BCs expressing the calcium biosensor GCaMP6f under the *ctbp2 (ribeyeA)* promoter fused to the synaptic protein Synaptophysin (Dreosti et al. 2009; Rosa et al. 2016). We focussed on BCs (Euler et al. 2014) as (i) they directly and differentially collect inputs from all photoreceptors to form the basis of colour vision (Li et al. 2012; Behrens, Schubert, Haverkamp, Euler & Berens 2016), (ii) they are the only neuron class that contacts all other neurons in the retina and (iii) they directly drive retinal ganglion cells, the eye’s connection to the brain. Individual pre-synaptic terminals of BCs can be resolved while imaging the entire depth of the inner retina (Dreosti et al. 2009) (Fig. 3a).

**Figure 3.**
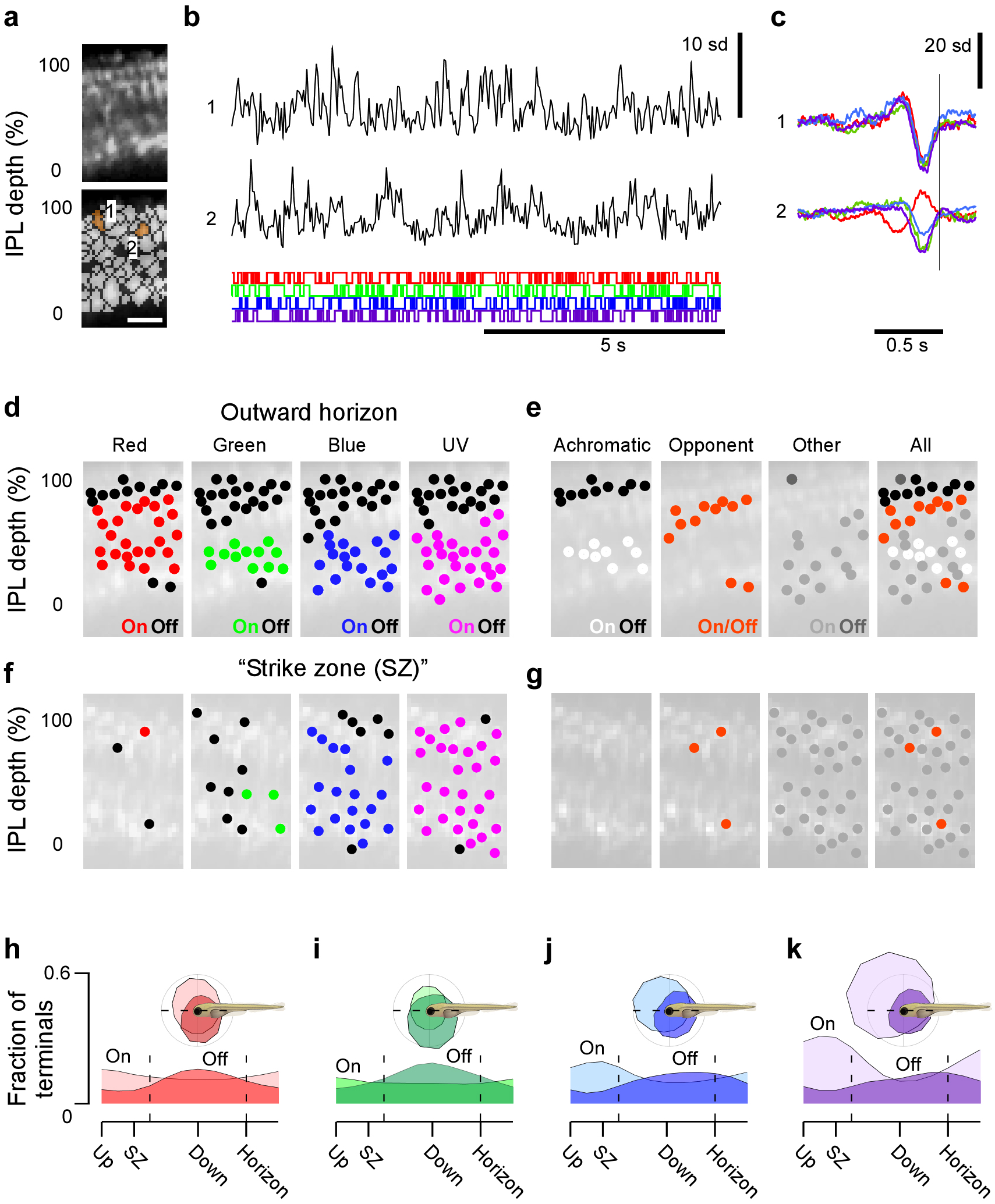
Surveying inner retinal chromatic responses *in vivo*. **a,** 2-photon scan-field (32 x 64 pixels, 15.625 Hz) of a nasal IPL section (outward horizon) in a *Tg(-1.8ctbp2:SyGCaMP6)* larvae for simultaneous recording of light-driven calcium responses across the entire IPL depth at single terminal resolution (top) and regions of interest (ROIs, bottom). Scalebar 5 μm. **b,** example of calcium responses to tetrachromatic binary white noise stimulation (12.8 Hz, Methods) of two ROIs highlighted in (a). **c,** tetrachromatic linear filters (“kernels”) recovered by reverse correlation of each ROI’s response with the noise stimulus (b). The colour code indicates the stimulus channel (R, G, B, U cf. SFig. 3a). **d,** For each stimulus channel, we classified each ROI’s kernel as either “On” (in red, green, blue or purple), “Off” (black), or non-responding (no marker) and plotted each response over the anatomical scan image (Methods). **e,** By comparison across the four stimulus channels, we then classified each ROI as either achromatic “Off” (R+G+B+U Off, black) or “On” (R+G+B+U On, white), Colour opponent (any opposite polarity responses in a single ROI, orange) or “other” (grey) and again plotted each ROI across the IPL to reveal clear chromatic and achromatic layering in this scan. **f,g,** As (d,e), but for a scan taken in the temporo-ventral retina which surveys the world in front of the animal just above the visual horizon. This zone is critical for prey capture, and was thus dubbed “strike zone”. **h-k,** Distribution of all On- and Off-responses per stimulus channel (h, red; i, green; j, blue; k, UV) based on n = 4,099 / 6,565 ROIs that passed a minimum quality criterion (Methods) sampled from across the entire sagittal plane (115 scans, 12 fish).

To estimate each BC terminal’s chromatic sensitivity, we used a tetrachromatic “noise” stimulus (Fig. 3b,c). Specifically, each of four LEDs that were spectrally matched to the absorption peaks of the four cone-opsins (SFig. 3a) were presented to the live larvae through the objective and flickered in a known random binary sequence at 12.8 Hz (Fig. 3b). Using reverse correlation (Chichilnisky 2001), we then recovered four temporal response kernels for each BC terminal (Franke et al. 2017), one for each LED and thus effective cone-type input (Fig. 3c). This revealed different chromatic sensitivities in different BC terminals. For example, some terminals displayed near-equal sensitivity to all four LEDs, indicating a wavelength-invariant response preference (achromatic terminals, Fig. 3a-c, ROI 1). Other terminals had kernels with opposite polarity across LEDs (colour opponent terminals, Fig. 3a-c, ROI 2). In an example scan from the “outward horizon” for all opsin channels, the majority of Off- and On-responses occurred in the upper and lower part of the IPL, respectively (Fig. 3d,e), in line with mammalian inner retinal circuits (Masland 2001; Wässle 2004; Euler et al. 2014; Franke et al. 2017). However, the transition depth between On- and Off-bands differed between cone-channels, with the R-channel transitioning closer to the inner nuclear layer (INL) than the other three. As a result, two achromatic bands at the top (RGBU_Off_, black) and in the centre (RGBU_On_, white) of the IPL were separated by a layer of R_On_/GBU_Off_ colour opponent responses (orange) (Fig. 3e). Additional R(G)_Off_/BU_On_ opponent responses occurred at the lower edge of the IPL. The remaining response types were classified as “other” and were mostly confined to the On-channel in the lower part of the IPL (grey). Accordingly, in this part of the eye the inner retina was organised into distinct functional layers. Moreover, as predicted from natural light, all colour opponent terminals computed short-versus long-wavelength chromatic contrasts. However, this functional layering was not consistently observed in other parts of the retina. In an example scan taken from the strike zone, nearly all terminals exhibited strong U(B)-On responses that reached far into the upper sublamina, while responses to R and G stimulation all but disappeared (Fig. 3f,g) – in striking agreement with the predicted need for dedicated UV-On prey-capture circuits in this part of the eye.

Together, these two examples demonstrate that the larval zebrafish IPL is functionally highly anisotropic. To systematically assess how BC responses are distributed across the eye, and which specific chromatic and colour opponent computations predominate, we recorded from a total of n=6,568 synaptic terminals across the sagittal plane (n = 115 scans, 12 fish at 7-8 *dpf*), out of which n=4,099 (62%) that passed a quality criterion (SFig. 3b,c, Methods) were used for further analysis. All recordings were taken in the same sagittal plane used for anatomy (cf. Fig. 2f). This dataset showed that the zebrafish larval retina is functionally highly anisotropic (Fig. 3h-k, cf. SFig. 3d). For example, independent of wavelength, On- and Off-responses were systematically biased to the upper- and lower-visual fields, respectively. Here, a disproportionate number of U_On_ responses surveyed the strike zone (Fig. 3k). What is the functional distribution of BCs across the larval zebrafish eye, and what do they encode?

## Large-scale functional anisotropies of the inner retina match natural spectral statistics

To assign BCs to functional clusters, we used a Mixture of Gaussian model to sort terminals independent of eye position based on their full temporo-chromatic response kernels (Fig. 4, Methods). BC terminals fell into n=26 clusters, which were further grouped into four major response groups: n=5 achromatic clusters (C_1-5_, Fig. 4a), n=9 UV(B)-monochromatic clusters (C_6-14_, Fig. 4b), n=6 chromatic clusters (C_15-20_, Fig. 4c), n=5 colour opponent clusters (C_21-25_, Fig. 4d) and n=1 discard cluster (C_x_, Fig. 4e, Methods). These groups were defined based on the relative amplitudes and polarities of each cluster mean’s four chromatic kernels (Methods): Equal polarity equal gain (achromatic), equal polarity different gain (chromatic and UV(B)-monochromatic) or different polarity (colour opponent). In addition, we distinguished UV(B)-monochromatic clusters from other chromatic clusters in view of the hypothesised behavioural relevance of such a channel (Fig. 1n). Their abundance and extreme short-wavelength bias indicate the existence of a dedicated UV-system that is not integrated with other chromatic retinal circuits.

**Figure 4.**
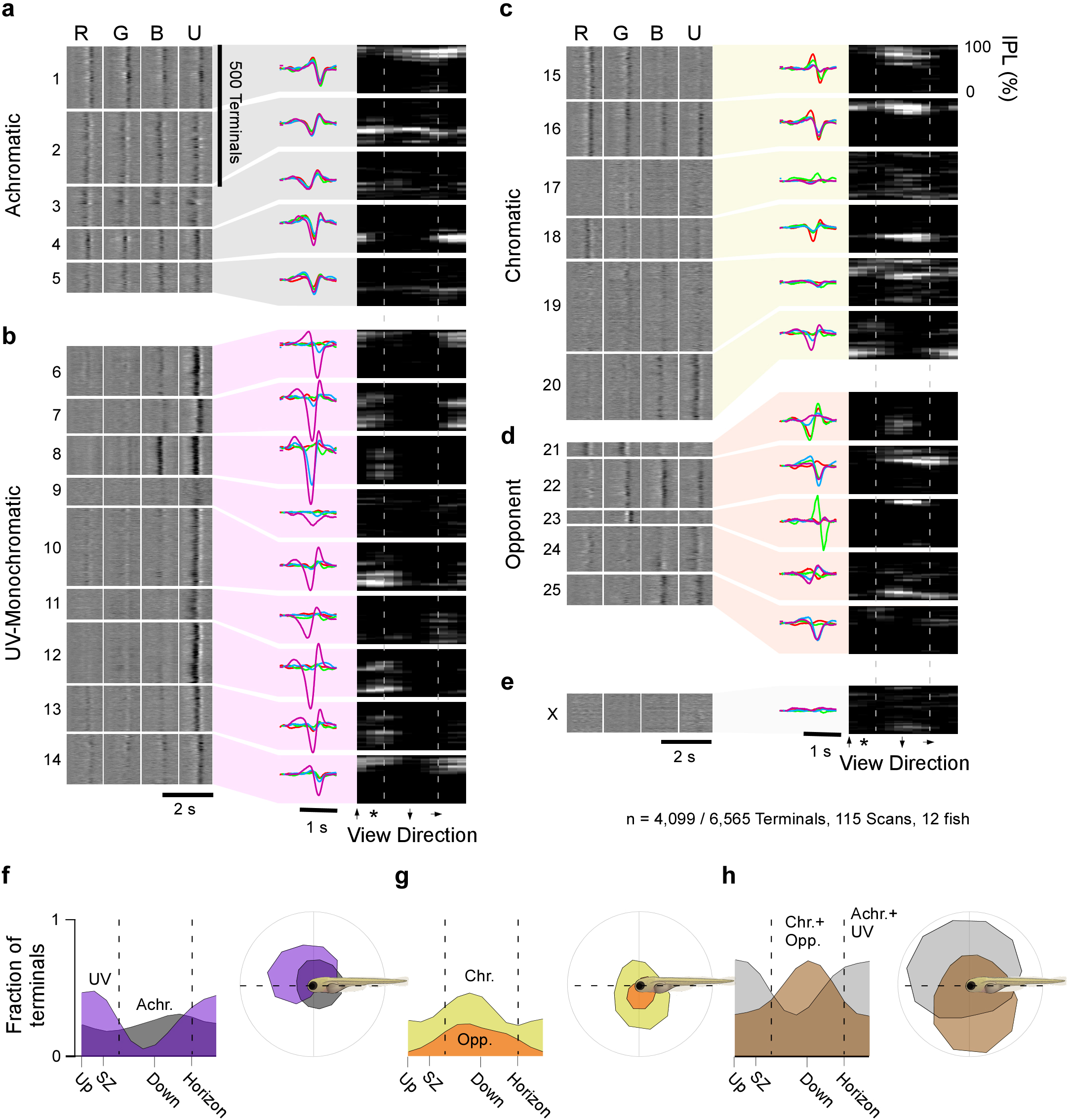
The functional organisation of the larval zebrafish eye. Mixture of Gaussian clustering of all n=4,099 responding terminals based on the full waveforms of their tetrachromatic kernels, with cluster number limited by the Bayesian Information Criterion (BIC), yielded 25 clusters (C_1-25_) and 1 discard cluster (C_x_). For simplicity, each cluster was further allocated to one of four major response groups (Methods): **a,** achromatic (C_1-5_), **b,** UV(B)-monochromatic (C_6-14_), **c,** chromatic (C_15-20_) and **d,** colour opponent (C_21-25_). **e,** Discard cluster C_x_. For each cluster, shown are the time-course of each kernel (left heatmaps, lighter shades indicating higher values), the cluster-means (middle) and their anatomical distribution across IPL depth (y axis) and position in the eye (x-axis, right heatmaps, lighter shades indicate higher abundance). Dashed lines indicate the forward and outwards horizon, the asterisk denotes the position of the strike zone. The height of each cluster’s left heatmap indicates its number of allocated terminals. **f-h,** Linear (left) and polar (right) histograms of terminal abundance of the functional groups defined in (a-e) across the larval zebrafish’s visual space. **f,** UV(B)-monochromatic (purple) and achromatic (grey) groups, **g,** chromatic (yellow) and colour opponent (orange) groups, **h,** summed UV(B)-monochromatic and achromatic groups (grey) versus chromatic and colour opponent groups (brown).

For each cluster, we computed the anatomical distribution across the eye and IPL depth (right insets). This revealed that no functional BC cluster, nor any major functional group (Fig. 4f-h), was uniformly distributed across the entire field of view. Instead, most clusters predominated in either the upper or lower visual field, with some clusters in addition exhibiting a secondary bias to predominately looking forwards or outwards. In agreement with our predictions, all UV(B)-monochromatic clusters were strongly biased to the upper and forward-facing visual field (Fig. 4b,f), while all colour opponent clusters were skewed towards the lower and outwards facing visual field (Fig. 4d,g). In fact, there were effectively no colour opponent terminals that survey the world directly upwards through the nearly achromatic Snell’s window. Together, all circuits potentially dealing with colour (all chromatic and opponent clusters) surveyed the lower and outward horizontal visual field, while all circuits likely to deal with achromatic computations (all UV(B)-monochromatic and achromatic clusters) were skewed towards the upper and frontal visual field (Fig. 4h). Moreover, four out of five colour opponent clusters computed short-versus long-wavelength chromatic antagonisms (reminiscent of PC2 from natural scenes), while the remaining single cluster (C_23_) compared G- to all other channels (reminiscent of a mix of PCs 3 and 4) (Fig. 4d, cf. Fig. 1k). This set of functional anisotropies of the larval zebrafish inner retinal circuitry is in strong agreement with the distribution of behaviourally meaningful chromatic content in nature (Fig. 1m,n).

## The functional layering of the larval zebrafish inner retina

Next, we assessed how different response types were distributed across the layers of the IPL (Fig. 5). Unlike in mammals (Euler et al. 2014), BCs in most non-mammalian vertebrates, including zebrafish, can have mono-, bi- and tri-stratified morphologies (Li et al. 2012; Connaughton & Nelson 2000). In agreement, terminals falling into individual clusters were mostly distributed across 1-3 major IPL bands (Fig. 4a-e, right insets), indicating that each could principally be linked to a single or small number of BC types. To establish the major trends in functional organisation of the IPL, we subdivided each of the four major response groups (achromatic, UV-monochromatic, chromatic, colour opponent) into their “On” and “Off”-dominated response-types and assessed the distribution of each of the resultant eight groups across the depth of the IPL for different positions in the eye (Fig. 5a).

**Figure 5.**
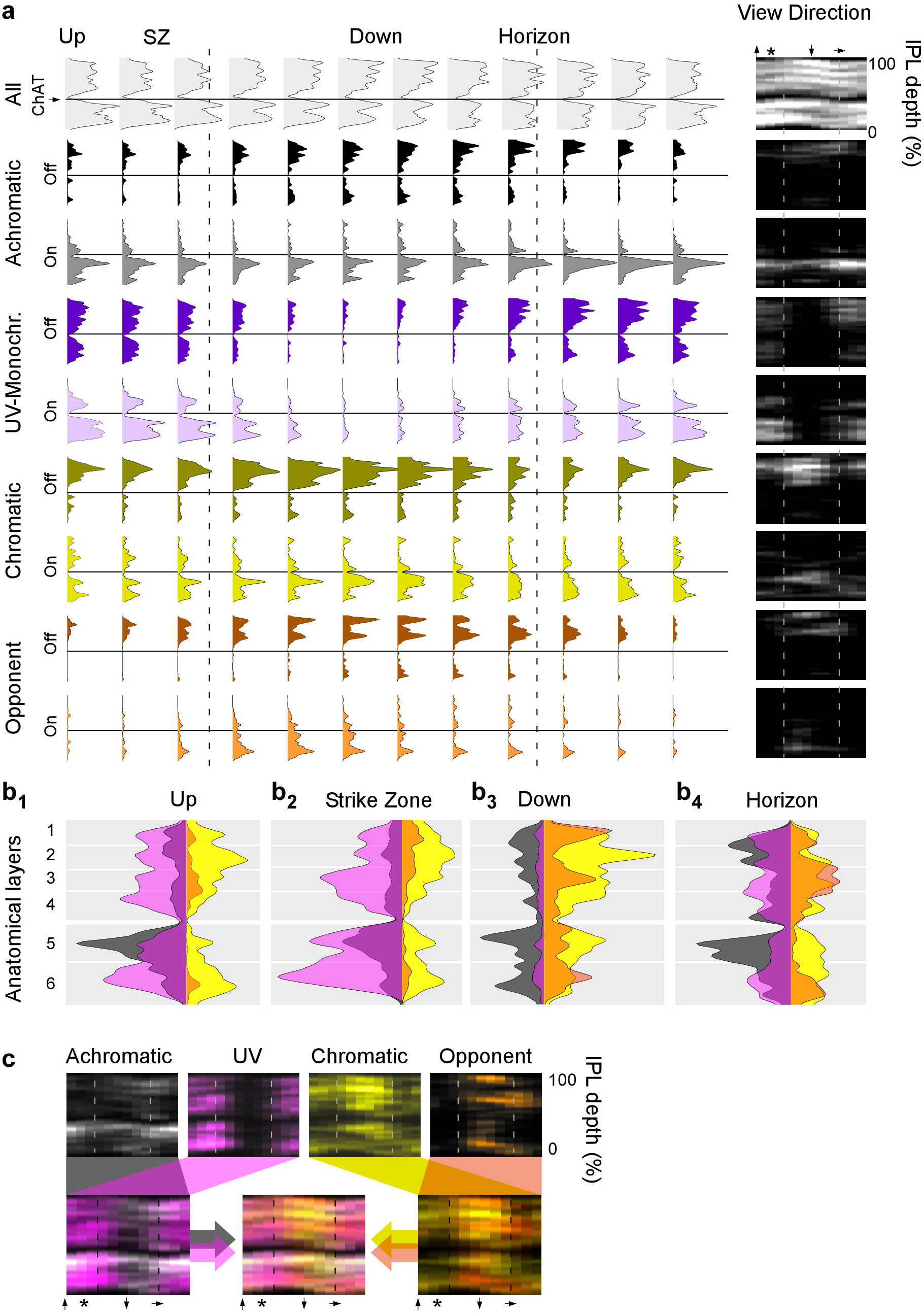
Distribution of function across the IPL. **a,** Histograms of terminal abundance across IPL depth (y axis) and position in the eye (set of histograms) for each functional group (cf. Fig. 7), divided by On- and Off-dominated responses as indicated. In addition, the distribution of all n = 6,565 scanned terminals independent of response-quality is plotted to reveal the anatomical distribution of all BC terminals (top, light grey). Heatmaps to the right show the same data in a single image. Asterisk denotes the position of the strike zone. Dashed lines indicate the forward and outward visual horizon. Solid horizontal lines indicate the position of the lower ChAT band as an anatomical reference. **b_1-4_,** On/Off-collapsed histograms of the four response groups for four example regions (up: ventral, strike zone: temporo-ventral, down: dorsal and outward horizon: nasal as indicated) summarise the functional IPL layering across eye positions, with approximate anatomical layers indicated in the background shading. For clarity, achromatic and UV(B)-monochromatic histograms are x-axis-reversed. **c,** Colour-coded response groups (top) plotted against eye position (x) and IPL depth (y) and merge (bottom). Throughout, colours indicate the functional groups: Achromatic (grey/black), UV(B)-monochromatic (purple/violet), chromatic (yellow/beige) and colour opponent (orange/brown).

**Figure 6.**
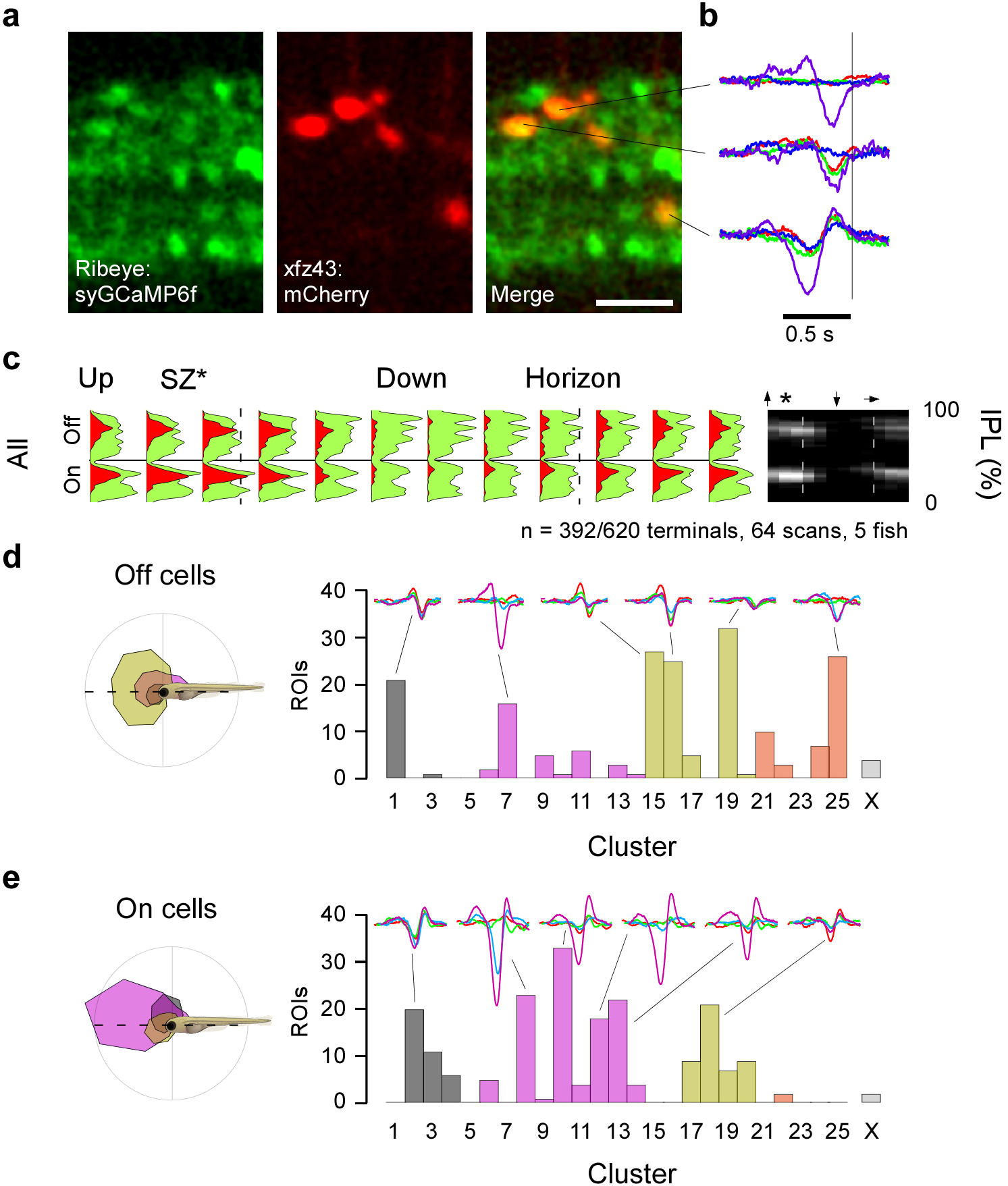
Distribution of genetically defined BC-types. **a,** High-resolution 2-photon scan of a ventro-nasal (up/outward) IPL section in 7 *dpf* larvae expressing SyGCaMP6f under *ctbp2* promoter (green) as well as mCherry under *xfz43* (red). Scalebar 5 μm. **b,** Subsequent higher rate scans during light-stimulation allowed recovering tetrachromatic kernels from individual xfz43-positive terminals as before (right). **c,** Distribution of 392 / 620 xfz43-positive BC terminals (64 scans, 5 fish) that passed our response criterion (red, Methods) across the IPL (y) and eye (x), superimposed on the distribution of all terminals from the same scans (green). The heatmap on the right shows only xfz43 positive terminals. Dashed lines indicate the forward and outward horizon, while the solid horizontal line indicates the position of the lower ChAT band. **d,** Allocation of all xfz43-positive anatomical Off terminals to functional clusters (right) and distribution of these terminals across the eye by functional group (left). **e,** As (d), but for xfz43-positive anatomical On-cells.

**Figure 7.**
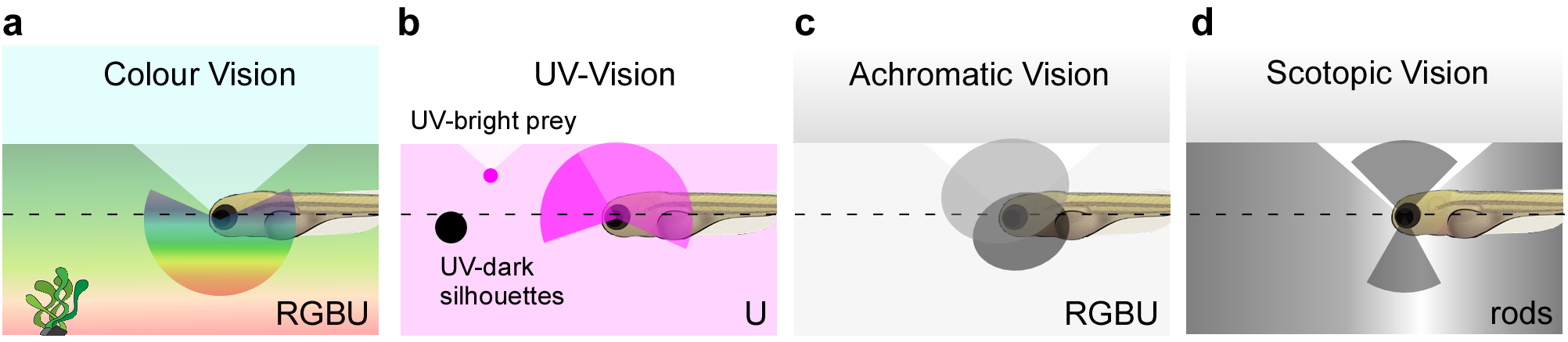
The larval zebrafish eye’s chromatic organisation for vision in nature. **a**, Circuits for colour vision are generally biased to the horizon and lower visual field where most chromatic content is found in nature. **b,** Circuits for UV(B)-monochromatic vision dominate the upper and frontal visual field and may be used for prey-capture and the detection of UV-dark silhouettes against a UV-bright background. **c,** Achromatic circuits are found throughout the eye, with On- and Off- circuits dominating the upper-frontal and loweroutward facing visual fields, respectively. **d,** Rod-circuits are exclusively used to survey the sky directly above and the ground reflection directly beneath the animal, where most photons can be caught. In each image, the triangular area above the animal depicts Snell’s window, and the visual horizon is indicated by a dashed line.

Most response groups (Fig. 5a, rows 2-9) were largely restricted to subsets of the larval zebrafish’s six anatomical IPL layers (row 1, black). However, some functions were distributed more broadly than others. For example, most achromatic On-terminals (row 3, grey) occurred just beneath the outer ChAT band (GCL side), and could be found in any eye-position – albeit at varying densities. In contrast, UV(B)-monochromatic On-terminals occurred across the entire outer part of the IPL (layers 3-6) but remained near exclusively restricted to the upper visual field (row 5, lilac). Other “functions” were tightly restricted in both visual field and IPL depth. For example, colour opponent Off-dominated terminals were near exclusively found in layers 1 and 3, and only in the lower visual field (row 8, brown). Next, we again combined “On” and “Off” versions of each response group for clarity and superimposed the resulting four histograms of the four response groups for different eye positions (Fig. 5b,c). Looking up, each IPL layer contained a substantial fraction of UV(B)-monochromatic terminals; only in layers 2 and 5 there were more chromatic and achromatic terminals, respectively (Fig. 5b_1_). In the strike zone, this UV(B)-dominance further intensified, and shifted towards the Off-bands in the lower IPL – in line with UV-On circuits aiding the detection of paramecia (Fig. 5b_2_, cf. Fig. 1n). In contrast, there were effectively no UV-monochromatic terminals looking down, and the IPL appeared more neatly divided into layers differentially encoding colour opponent, chromatic and achromatic information (Fig. 5b_3_). Finally, IPL circuits surveying the outward horizon had approximately balanced numbers of terminals from each functional group, and a similarly neat functional organisation with IPL depth as observed for the lower visual field (Fig. 5b_4_).

The complexity of functional layers differed markedly, in particular between the strike zone and upper visual field (Fig. 5b_1,2_), compared to the lower visual field and outward horizon (Fig. 5b_3,4_). In the latter, the total number of peaks often used as a tell-tale for individual IPL strata far exceeded the 6 traditionally used anatomical layers of the larval zebrafish IPL. This suggests that this part of the eye harbours more diverse BC circuits than what is required in much more “simple”-appearing circuits surveying the upper visual field.

A neat division of function into specific layers of the IPL for surveying the ground and outward horizon, though novel in its chromatic scope and complexity, is consistent with current knowledge on the functional organisation of the vertebrate inner retina (Euler et al. 2014; Masland 2001; Wässle 2004; Franke et al. 2017). However, the overrepresentation of the UV(B)-channel in the upper and frontal visual fields, despite the presence of all other cone types, is striking. Here, most visual functions appear to draw near exclusively on On- and Off-UV(B)-monochromatic channels at the expense of both colour vision and the Off achromatic channel. How does the eye build this rich functional division across visual space?

## Building a functionally anisotropic retina

Across all vertebrate retinas studied to date there are distinct types of BCs, each with a unique anatomy, synaptic connections and functional properties (reviewed in (Euler et al. 2014)). Both larval and adult zebrafish have ~20 morphologically distinct BC types, with a broad diversity of chromatic connections to cones in the outer retina (Li et al. 2012; Connaughton & Nelson 2000; Vitorino et al. 2009), many more than the dichromatic mouse (Behrens, Schubert, Haverkamp, Euler, Haverkamp, et al. 2016; Helmstaedter et al. 2013; Kim et al. 2014; Greene et al. 2016; Wässle et al. 2009). Two non-mutually exclusive design strategies may underlie the observed set of functional anisotropies; First, different types of BCs with different functions might specifically exist only in certain parts of the retina (Zhang et al. 2012; Bleckert et al. 2014). This hypothesis is, for example, supported by the absence of the characteristic large terminals of On-type mixed BCs outside the ventral- and dorsalmost retina (Fig. 2h,j), where rods are located (Fig. 2b,c). Second, the same types of BCs may exist across the entire retina, but shifting function with position (Baden et al. 2013; Joesch & Meister 2016; Chang et al. 2013; Sabbah et al. 2017).

We set out to test these two possibilities experimentally. For this, we used the xfz43 marker line, which labels at least three morphologically well-characterised types of BCs with different, known anatomical connections in the outer retina (D’Orazi et al. 2016). One Off- and one On-stratifying xfz43-positive BC type each preferentially contacts R- and G-cones across their large dendritic fields. Both are thus predicted to exhibit a RG-biased physiology. A third, smaller On-stratifying xfz43 type indiscriminately samples from all cones, and is therefore expected to encode On-achromatic features.

To selectively record from these cells’ synaptic terminals, we crossed our *Tg(-1.8ctbp2:SyGCaMP6)* line to *Tg(xfz43:Gal4;UAS:ntr-mCherry)* animals (Fig. 6a,b). As before, this allowed recording from all BC terminals, but in addition labelled xfz43-cells are simultaneously recorded in the red fluorescence channel. This dataset (64 scans, 5 fish) represents a subset of the complete BC-survey presented previously (Figs. 3-5) and consisted of n=620 xfz43-positive terminals, of which the 392 (63%) that passed our quality criterion (Methods) fell into 2 main IPL bands (Fig. 6c, red).

Next, we functionally assessed xfz43 Off- and On-stratifying terminals separately. In agreement with their stratification, Off- and On-stratifying terminals fell into functional Off-and On-clusters, respectively (Fig. 6d,e). However, the presumed single xfz43 Off-type (D’Orazi et al. 2016) fell into 6+ functional clusters that spanned all four major functional groups (Fig. 6d). Similarly, the presumed two On-type xfz43 cells were sorted into 6+ clusters that spanned 3 major functional groups (Fig. 6e). In fact, individual examples clearly demonstrated this functional diversity within a genetic type, irrespective of any clustering (not shown). This extreme functional diversity of all three presumed “clean”, genetically defined BC types strongly suggests that - in the larval zebrafish eye - genetics alone do not predict a BC’s physiology. Instead, a cell’s functional identity appears to fundamentally depend on its surrounding neuronal network in different parts of the eye.

However, the allocation of xfz43-cells to functional clusters was far from random. For example, xfz43 Off-terminal allocated clusters C_1_, C_15_ and C_17_ all exhibited at least a small RG-bias, consistent with these cells’ known connectivity in the outer retina (D’Orazi et al. 2016). Similarly, cluster C_18_, which captured most chromatic xfz43 On-terminals, had an RG-biased physiology, while the largely achromatic cluster C2 might reflect the cone-unselective xfz43 cells. In each case, this mainly leaves several UV(B)-dominated response clusters that are not explained by these cells’ cone-selectivity (C_6-8,12,13_).

Are these UV-clusters generated by UV-cone inputs with unusually high gain? For example, the small On-type xfz43 cell indiscriminately integrates the outputs of fewer than 10 cones (D’Orazi et al. 2016). Here, a hypothetical high-gain UV-input from only two or three cones could bias a cell towards a UV-dominated response. While this hypothesis clearly needs further exploration, already here several lines of evidence point at this as one mechanism of functional diversification across the larval zebrafish IPL. First, under natural light, UV-cones receive ~15 times fewer photons than red cones (Fig. 1h), prompting the need for a high-gain amplification system for short-wavelength visual processing. In agreement, in mice the gain of UV-cones appears to be higher than that of M-cones (Breuninger et al. 2011; Baden et al. 2013). Second, UV-cones numerically dominate the frontal- and upper visual field (Fig. 2b-d). Third, UV-responses overall dominate the IPL in this part of the eye (Figs. 3k, 4b,f, 5), with their kernel amplitudes often exceeding those of any other opsin channels more than 10-fold, despite the presence of all other cone types. Taken together, it therefore seems likely that at least to some extent larval zebrafish BC types with specific function exist in only parts of the eye, but that in addition more large-scale outer- and/or inner-retinal circuits can “override” this genetically defined functional organisation.

## Conclusion and outlook

We have shown that inner retinal circuits of larval zebrafish are exquisitely matched to their natural visual environment on several levels. First, chromatic circuits are systematically integrated by a neatly layered inner retina, but only at the horizon and the lower visual field, which in nature contain the most chromatic information (Fig. 7a, cf. Figs. 1e,k, 4g,h, 5b). Here, the chromatic computations performed by these circuits match the differential predominance of different natural chromatic contrasts and behavioural demands (Fig. 4d, Fig. 1k). The upper and frontal visual fields are dominated by UV-driven circuits, a specialisation that begins with an anisotropic arrangement of UV-cones across the eye (Fig. 2b-d) and is mirrored across the ventral inner retina, apparently at the expense of circuits serving colour vision and near inner retinal organisation (Fig. 7b, cf. Figs. 3k, 4f, 5). This UV-dominance is likely linked to the need to spot nearby UV-bright microorganisms (Novales Flamarique 2012) as well as the ability to detect UV-dark objects on the backdrop of further UV-bright organic matter dissolved in the water (Losey et al. 1999; Nava et al. 2011; Goldsmith 1994). Achromatic cone-driven circuits are used across the entire visual field, but differentially use On- and Off dominant regions in the upper and lower visual field respectively, possibly to drive the dorsal righting response which helps fish maintain an upright posture by aligning their vertical body axis with the brightness gradient of light from the sky (Fig. 7c, cf. Fig. 3h-k). Finally, rod-driven circuits exclusively survey the visual field above and below the animal, likely to capitalise on the additional light caught through Snell’s window and its ground reflections (Fig. 7d, cf. Fig. 2b, right). To what extent this set of striking regional specialisation of the larval zebrafish visual system is already established in the outer retina, how it is reflected in the adult (Pita et al. 2015), and how it is used in the retinal output and brain to ultimately drive behaviour will be important to address in future studies.

## Methods

No statistical methods were used to predetermine sample size.

### Field sites

Six field sites were visited in West Bengal, India (Supplementary Fig. 1) in May 2017, just before the monsoon season. The global positioning coordinates of each site were: Site 1 (lat. 26.531390, long. 88.669265), site 2 (lat. 26.528117, long. 88.641474), site 3 (lat. 26.841041, long. 88.828882), site 4 (lat. 26.792305, long. 88.588003), site 5 (lat. 26.903202, long. 88.554333) and site 6 (lat. 26.533690, long. 88.648729). Zebrafish of all ages were found mostly in shallow pools of water adjacent to bigger streams (with exception of one deeper fish farm pond, site 6), in agreement with previous work (Arunachalam et al. 2013; Spence et al. 2008; Parichy 2015). The visited sites varied substantially with the type of habitat (different sized streams, stagnant fish farm pond), the amount of vegetation above and under water, the type of substrate and the current of the water (Supplementary Data). For analysis, all recorded data was combined without prior selection.

### Hyperspectral imaging

To gather hyperspectral images, we used a custom made, waterproofed hyperspectral scanner (Nevala et al. 2017) built around a commercial spectrometer (Thorlabs CCS200/M, 200-1,000 nm). In short, two spectrally broad mirrors mounted on top of servo-motors were controlled by an Arduino Uno microcontroller to iteratively bring different positions in visual space into the active part of the spectrometer to build up a complete hyperspectral image (Baden et al. 2013). 1,000 Scan-points were spaced 1.6° and defined by a Fermat’s spiral, followed by a custom path-shortening algorithm. Spectra were recorded using the spectrometer software OSA (Thorlabs). We used diving-weights to stabilise the scanner under water. In addition, the scanner-case was placed inside a hard-plastic box to maintain the upright position with a UV-transparent window pointing forward.

After placing the scanner to its <50 cm depth underwater position, we waited up to 5 minutes for any stirred-up debris to settle. All n=31 scans were taken during the day between 11am and 5pm; the weather conditions varied from slightly cloudy to clear sky, but remained constant for individual measurements. Time for one scan acquisition varied between 4 and 8 minutes, depending on the set mirror-move times (200-500 ms) and integration times (80-200 ms) which were adjusted for each measurement to yield an approximately consistent signal-to-noise independent of absolute light intensity in each scene. Finally, in each case in addition to the scan a 180° still image was taken approximately at the scanner position with an action camera (Campark ACT80 3K 360°). Stills were mapped to the 2D plane by a standard angular fisheye projection to 720x720 pixels (0.25° per pixel).

### Animals and tissue preparation

All procedures were performed in accordance with the UK Animals (Scientific Procedures) act 1986 and approved by the animal welfare committee of the University of Sussex. For all experiments, we used 7-8 days post fertilisation (*dpf*) zebrafish (*Danio rerio*) larvae of either sex. The following transgenic lines were used: *Tg(-1.8ctbp2:SyGCaMP6)*, *Tg(xfz43:Gal4;UAS:ntr-mCherry;-1.8ctbp2:SyGCaMP6)* (Zhao et al. 2009), *Tg(opn1sw1:GFP)* (Takechi et al. 2003), *Tg(-3.2opn1sw2:mCherry)* (Salbreux et al. 2012), *Tg(thrb:Tomato)* (Suzuki et al. 2013). In addition, a *Tg(xops:ntr-mCherry)* line was generated by injecting pXops-nfsB-mCherry plasmid into one-cell stage eggs and subsequently screening for the expression of mCherry among progenies of injected fish. pXops-nfsB-mCherry plasmid was constructed by replacing EGFP with nfsB-mCherry in XopsEGFP-N1 plasmid (Fadool 2003).

Owing to the exploratory nature of our study, we did not use randomisation and blinding. Animals were housed under a standard 14:10 day/night rhythm and fed 3 times a day. Animals were grown in 200 μM 1-phenyl-2-thiourea (Sigma) from 1 *dpf* to prevent melanogenesis (Karlsson et al. 2001). For 2-photon *in-vivo* imaging, zebrafish larvae were immobilised in 2% low melting point agarose (Fisher Scientific, Cat: BP1360-100), placed on the side on a glass coverslip and submersed in fish water. Eye movements were further prevented by injection of α-bungarotoxin (1 nl of 2 mg/ml; Tocris, Cat: 2133) into the ocular muscles behind the eye. For immunohistochemistry, larvae were culled by tricaine overdose (800 mg/l) at 7-8 *dpf*. Whole larvae were fixed in 4% paraformaldehyde for 25 min before being washed in phosphate-buffered saline (PBS).

### Two-photon Ca^2^+ imaging and light stimulation

We used a MOM-type two-photon microscope (designed by W. Denk, MPI, Martinsried; purchased through Sutter Instruments/Science Products). Design and procedures were described previously (Euler et al. 2009). In brief, the system was equipped with a mode-locked Ti:Sapphire laser (Chameleon Vision-S, Coherent) tuned to 927 nm, two fluorescence detection channels for GCaMP6f (F48x573, AHF/Chroma) and mCherry (F39x628, AHF/Chroma), and a water immersion objective (W Plan-Apochromat 20x/1,0 DIC M27, Zeiss). For imaging mCherry, we used 960 nm excitation instead. For image acquisition, we used custom-written software (ScanM, by M. Mueller, MPI, Martinsried and T. Euler, CIN, Tuebingen) running under IGOR pro 6.3 for Windows (Wavemetrics), taking 64 × 32 pixel image sequences (15.625 frames per s) for activity scans or 512 × 512 pixel images for high-resolution morphology scans.

For light stimulation, we focussed a custom-built stimulator through the objective, fitted with band-pass-filtered light-emitting diodes (LEDs) (‘red’ 588 nm, B5B-434-TY, 13.5cd, 8°, 20 mA; ‘green’ 477 nm, RLS-5B475-S; 3-4 cd, 15°, 20mA; ‘blue’ 415 nm, VL415-5-15; 10-16 mW, 15°, 20 mA; ‘ultraviolet, UV’ 365 nm, LED365-06Z; 5.5 mW, 4°, 20 mA, Roithner, Germany). LEDs were filtered and combined using FF01-370/36, T450/pxr, ET420/40m, T400LP, ET480/40x, H560LPXR (AHF/Chroma). The final spectra approximated the peak spectral sensitivity of zebrafish R-, G-, B-, and UV-opsins, respectively, while avoiding the microscope’s two detection bands (SFig. 3a). LEDs were synchronised with the scan retrace at 500 Hz. Stimulator intensity was calibrated (in photons per second per cone) such that each LED would stimulate its respective zebrafish cone-type with an equal number of photons (~10^5^ photons per cone per s). Assuming an effective absorption coefficient of ~0.1, this translates to ~10^4^ photoisomerisations per cone per s (R*), a low photopic regime. We did not attempt to compensate for cross-activation of other cones, and relative LED-versus-opsin cross sections are listed in SFig. 3a. Owing to two-photon excitation of photopigments, an additional, steady illumination of ~10^4^ R* was present during the recordings (for detailed discussion, see (Baden et al. 2013; Euler et al. 2009)). For all experiments, the animal was kept at constant illumination for at least 5 seconds after the laser scanning started before light stimuli were presented. The only stimulus used throughout this work was a “tetrachromatic binary noise” stimulus. Here, each of the 4 LEDs was simultaneously but independently driven in a known binary sequence at 12.8 Hz for 258 seconds.

### Immunohistochemistry

For the IPL structural analysis, whole fixed larvae (7-8 *dpf*) were incubated in permeabilisation/blocking buffer (PBS with 0.5% Triton-X 100 and 5% normal donkey serum) for at least 10 min followed by 3-5 days incubation at 4°C with primary antibodies (chicken anti-GFP (AbCam, 13970, 1:500), goat anti-ChAT (Chemicon, AB144P, 1:50), rabbit anti-PKCα (Sigma, P4334, 1:100)). Samples were rinsed three times in phosphate buffered saline with 0.5% Trion-X 100 and incubated for another day with secondary antibodies and Hoechst 33342 (1:5000) for nucleus staining in permeabilisation/blocking buffer. Finally, samples were washed in PBS with 0.5% Triton-X 100 and mounted in mounting media (VectaShield, Vector, H-1000) for fluorescent imaging. Secondary antibodies used were as follows: Donkey anti-chicken IgG CF488A conjugate (Sigma, 1:500), Donkey anti-rabbit IgG CF568 conjugate (Sigma, 1:500), Donkey anti-goat IgG DyLight650 conjugate (BETHYL, 1:200). For photoreceptors, whole eyes were dissected from the animal at 7-8 *dpf* and subsequent tissues were subjected to immunohistochemistry as described above. Antibodies used are, a primary antibody: zpr-1 (ZFIN, 1:50), and a secondary antibody: Donkey anti-mouse IgG DyLight649 conjugate (Jackson Immunoresearch laboratories, 1:500). Confocal images were taken on Leica TCS SP8 or Olympus FV1000 using objectives 63x (HC PL APO oil CS2, Leica), 20x (HC PL APO Dry CS2, Leica), 60x (UPLSAPO oil, Olympus) or 20x (UPLSAPO oil, Olympus) at xy: 0.1-0.07 μm/pixel, and z-step: 0.25-0.3 μm for high-resolution images and 0.7-0.5 μm/pixel, and z-step: 2 μm for low magnification images. Images were median-filtered and contrast and brightness were adjusted in Fiji (NIH).

### Photoreceptor densities

Confocal image stacks of whole eyes were converted to spherical max-projection image stacks using custom-written scripts in IGOR pro 6.3 (Wavemetrics). The image plane crossing photoreceptor somata in this spherical projection was used to automatically identify cells using a threshold function in SARFIA (Dorostkar et al. 2010) running under IGOR Pro. After manual verification and correction in Fiji, photoreceptor positions were projected back onto a 3D hemi-sphere and density was calculated using custom IGOR Pro scripts. Density maps (Fig. 2a,b) are top-down views of these 3D hemispheres. For extracting density profiles linearlised against eye position (Figs. 2c,d) we computed the mean density in a ring between 112°-145° eccentricity, whose angular centre of mass corresponds to 130°. This “3D ring” was chosen as it corresponds to the same sagittal plane as surveyed for inner retinal anatomy and function (Fig. 2e, SFig. 2).

### Data analysis

Data analysis was performed using IGOR Pro 6.3 (Wavemetrics), Fiji (NIH) and Python 3.5 (Anaconda distribution, scikit-learn 0.18.1, scipy 0.19.0 and pandas 0.20.1).

### Pre-processing and receptive field mapping

Regions of interest (ROIs), corresponding to individual presynaptic terminals of BCs were defined semi-automatically by custom software (D. Velychko, cf. (Baden et al. 2016)). Next, the Ca^2+^ traces for each ROI were extracted and de-trended by high-pass filtering above ~0.1 Hz and followed by z-normalisation based on the time interval 1-6 seconds at the beginning of recordings using custom-written routines under IGOR Pro. A stimulus time marker embedded in the recording data served to align the Ca^2+^ traces relative to the visual stimulus with a temporal precision of 2 ms. We then mapped linear receptive fields of each ROI by computing the Ca^2+^ transient-triggered-average. To this end, we resampled the time-derivative of each trace to match the stimulus-alignment rate of 500 Hz and used thresholding above 0.7 standard deviations relative to the baseline noise to the times *t_i_* at which Calcium transients occurred. We then computed the Ca^2+^ transient-triggered average stimulus, weighting each sample by the steepness of the transient:

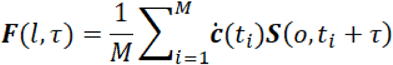

Here, *S*(*l*, *t*) is the stimulus (“LED” and “time”), *τ* is the time lag (ranging from approx. −1,000 to 350 ms) and *M* is the number of Ca^2+^ events. RFs are shown in z-scores for each LED, normalised to the first 50 ms of the time-lag. To select ROIs with a non-random temporal kernel, we first concatenated all four colour kernels to a single vector (X by 1) and computed the standard deviation across this vector. We used all ROIs with a standard deviation of at least two. The precise choice of this quality criterion does not have a major effect on the results.

### Feature extraction and Clustering

For each ROI, we concatenated the kernels for all colours, forming one 2,496-dimensional vector (4 times 649) per ROI. We then denoised this vector by using the reconstruction obtained from projecting it on the first 40 PCA components, capturing ~90% of the variance. We then followed a feature extraction and clustering pipeline described previously (Baden et al. 2016). We computed three PCA features on each colour channel individually, yielding a total of 12 features. They captured between 70 and 83% of the variance per channel. We fit a Gaussian Mixture Model the data, optimizing the Bayesian Information Criterion (BIC) for the number of mixture components.

The covariance matrix of each cluster was chosen to be diagonal and a regularisation term of 10^−6^ was added to the diagonal. The BIC curve was shallow between 22 and 27 clusters, with a minimum at 26. Spherical covariance matrices or the same covariance matrix for each cluster yielded higher BIC scores. Full covariance matrices yielded somewhat lower BIC scores with an optimum at a cluster number below 10. In this case, functionally heterogenous clusters were grouped together. This analysis was performed in Python 3.5 using scikit-learn implementations.

### Grouping of clusters into response groups

Each cluster was allocated into one of four response groups (n=25) or discarded (n=1). For each cluster mean and each channel, we first calculated the peak to peak amplitude in z-scores relative to each channels baseline, defined as the first 50 ms of each kernel. If the mean difference of the mode of all amplitudes between the UV and all other channels exceeded 35, that cluster was classified as UV(B) monochromatic (C_6-14_). Similarly, a single cluster with mean mode amplitude below 2 was discarded (C_x_). Next, we calculated the correlation between all pairs of channels as well as the variance between amplitudes, with the mean between amplitudes normalised to 1. If the mean correlation between all pairs exceeded 0.8 (i.e. similar waveforms) and the variance of amplitudes was below 0.09 (i.e. similar amplitudes), that cluster was classified as achromatic (C_1−5_). Finally, to distinguish remaining chromatic (C_15−20_) and colour opponent clusters (C_21− 25_), we also computed the mean of the mode of all correlations. If the mean of correlation equalled the mean of the mode of correlations (i.e. all kernels had the same polarity), that cluster was classified as chromatic. All remaining clusters were classified as colour opponent. Following this automatic pre-sorting, we manually reallocated three clusters that were misclassified due to low amplitudes of individual kernels: C_17_ and C_20_ were moved from colour opponent to chromatic as the very low amplitudes of the R-channel led to these clusters’ erroneous classification, and C_9_ was moved from the chromatic to the UV(B) monochromatic group as this cluster effectively only responded to UV-stimulation but the overall low-amplitudes led its misclassification. Finally, we also moved C_21_ from the chromatic to the opponent group. Here, the pronounced time-course difference between UV(B) and RG that leads to a clear opponency in the early kernel period was not picked up by our automatic sorting rules.

### Histograms against eye position

All histograms against eye position were smoothed using a circular 60° binomial (Gaussian) filter along the x-dimension and IPL depth histograms were in addition smoothed by a 5%-width Gaussian filter across the y-dimension. Moreover, all 2D histograms of both eye position and IPL depth (Figures 4-6) were warped to horizontally align the peaks of the major anatomical IPL layers across eye position (Fig. 5a, top row). Specifically, the IPL was compressed from the top by 5% at the outwards horizon and by 5% from the bottom of the IPL at the forward horizon, where the IPL is thickest (cf. Fig. 2k).

## SUPPLEMENTARY MATERIALS

### The cone-opsin complement of zebrafish

While larval (and adult) zebrafish have four cone-types and one rod-type, each cone’s *in vivo* action spectrum depends on several factors, including which opsin gene(s) are expressed, which retinal chromophore(s) are used, and what fraction of the spectrum of light is filtered out by the optical apparatus. To justify the choice of opsin templates used (cf. SFig. 3a) we will discuss each in turn.

Zebrafish have UV, blue, green and red sensitive cones in their retina. The maximal absorbance (λ_max_) for the UV sensitive cones (UVS) lies around 360-365 nm while the λ_max_ for the blue sensitive cones (SWS, short wavelength sensitive cone) is at 411nm (Allison et al. 2004; Hunt et al. 2001). For green and red cones (medium and long wavelength sensitive cones, MWS and LWS) the situation is more complex since these cones can express different types of opsins. Zebrafish use four MWS-cone opsins (RH2-1, RH2-2, RH2-3 and RH2-4) and two LWS-cone opsins (LWS1 and LWS2) (Chinen et al. 2003). All these opsins have different spectral sensitivities, and all are expressed in the retina (Takechi & Kawamura 2005). This variation is expected in a small spectral sensitivity shift in these cones with retinal position. Based on the abundance of the opsin type across the retina for the green and red cones during the early development of zebrafish larvae, we chose the most abundant opsin type in each case: the RH2-1 gene (λ_max_ at 467 nm) for the green cones and the LWS1 gene (λ_max_ at 548 nm) for the red cones.

In addition, vertebrates can use two different chromophores: 11-cis-retinal (vitamin A1) and 11-cis-3,4-didehydroretinal (vitamin A2). The A2 chromophore holds one extra double bond compared to A1, which lowers the energy needed to initiate the phototransduction cascade (Enright et al. 2015; Koskelainen et al. 2000; Loew & Dartnall 1976). By changing the chromophore from A1 to A2, a visual pigment’s peak spectral sensitivity can be shifted to longer wavelengths. While for UVS- and SWS-opsins, this switch has little effect, MWS-opsins can be shifted by 20 nm, while LWS-opsins shift up to 60 nm (Allison et al., 2004). This change can be triggered in adult zebrafish when treated with thyroid hormone (Allison et al., 2004; Enright et al., 2015), but there is no clear evidence for the zebrafish of any age holding the A2 chromophore under normal conditions. We therefore assumed that only the A1 chromophore is present. As such, λ_max_ values for the cone templates were set to 365 nm, 411 nm, 467 nm and 548 nm for the four cone types, respectively.

Other possible structures in the eye filtering the light reaching the light sensitive cells are the cornea, the lens and the vitreous body. All these parts can hold patches of pigments that mostly absorb shorter wavelengths (UV) affecting the spectrum of light that eventually reaches the photoreceptors (Douglas & McGuigan 1989). Whether or not the animal has these pigments varies widely across different species (Siebeck & Marshall 2001), but measurements are still lacking for the larval zebrafish. Notably, here the small size of the eye, and the fact that UV-cones exist throughout the retina, strongly suggest that UV-filtering by the optical apparatus in these animals is negligible. As such, we did not correct our opsin templates for any putative spectral related to the optical apparatus of the eye.

### The number of neurons in the 7-8 *dpf* larval zebrafish

Our photoreceptor-labelling experiments (Fig. 2) revealed that at 7-8 *dpf*, a single eye comprises approximately 10,000 photoreceptors (all cones and rods combined). From here, we then estimated HC and BC numbers as ~1,000 and ~25,000, assuming these neuron classes comprise 4 (HC) and 25 (BC) types that each tile the retina with no overlap, with each type on average contacting 30 (HC) and 10 (BC) PRs (D’Orazi et al. 2016; Li et al. 2009; Vitorino et al. 2009; Song et al. 2008). Finally, there are ~4,000 RGCs in the larval zebrafish eye (Robles et al. 2014), and from here we assumed that ACs make up at least another 4,000 neurons. This puts the total number of neurons for both eye added up ~88,000. In comparison to ~85,000-100,000 neurons in the brain excluding the eyes (Naumann et al. 2010), this estimate implies that about half the brain’s neurons are located in the eyes.

### Choice of age of zebrafish larvae

Throughout this work we used 7-8 *dpf* zebrafish larvae of either sex. At this age, zebrafish brain and retina conform to the anatomical structures of vertebrate nervous system, such as existence of canonical cell types and specific neural circuits (McLean & Fetcho 2011; Hoon et al. 2014). Supported by their visual circuits, 7-8 *dpf* zebrafish larvae perform a wide range of behaviours including reflexive responses as well as prey-capture and predator avoidance (Orger 2016; Friedrich et al. 2010), allowing them to feed independently, navigate their environment and avoid predators. Though under constant development, larval zebrafish of this age are therefore fully autonomous and highly visual animals. They have been used extensively to study vertebrate nervous system organisation and function including benchmark studies of whole-brain functional imaging (Keller et al. 2014) and serial-section electron microscopy (Hildebrand et al. 2017).

## SUPPLEMENTARY FIGURE LEGENDS

**Supplementary Figure 1 | related to Figure 1:**
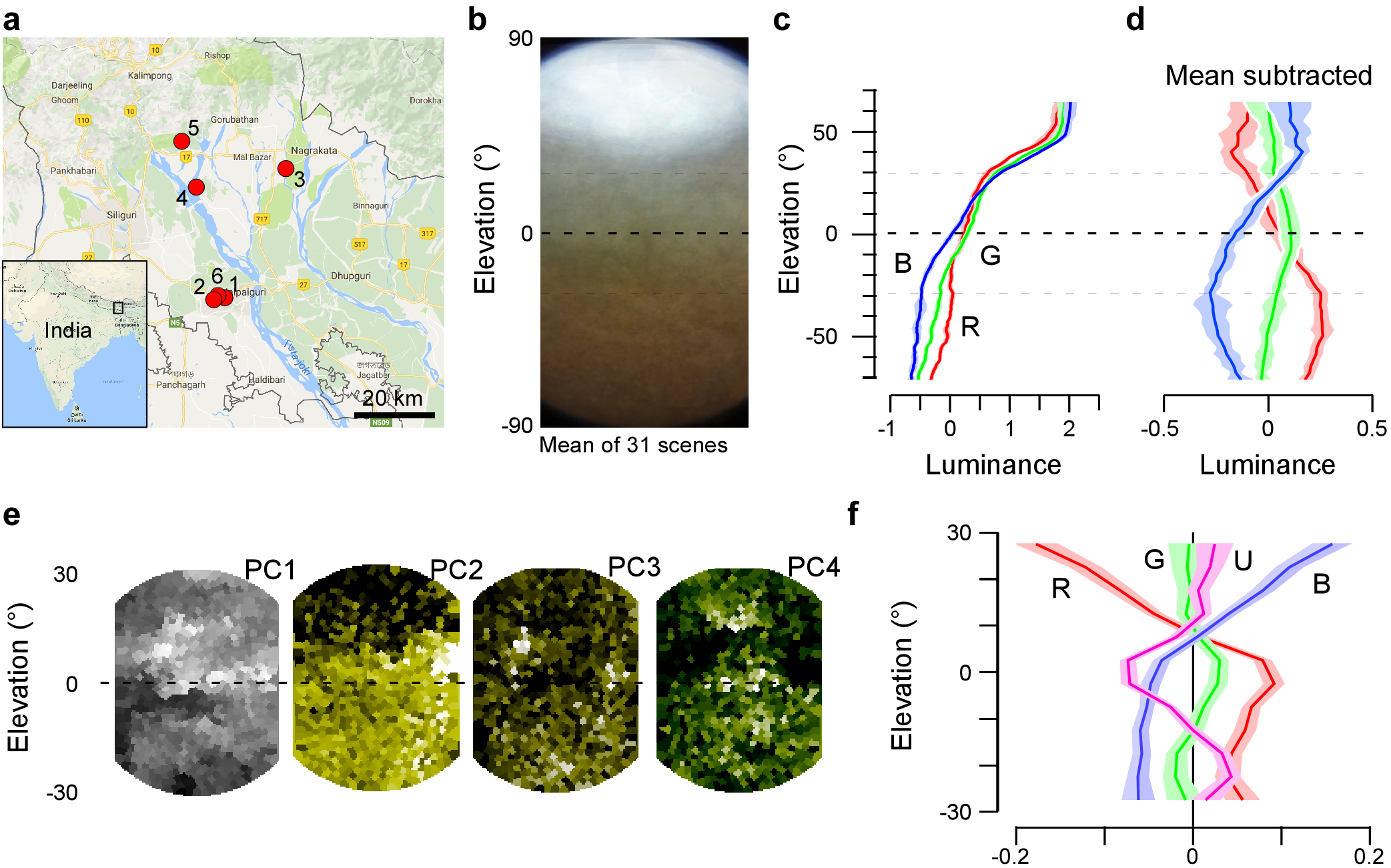
Distribution of chromatic content in the zebrafish natural visual world. **a**, Location of field sites visited in West Bengal, India. **b,** Mean of n=31 action camera images and **c,** mean z-normalised brightness of the red, green and blue camera channels across these scenes. **d,** As (c), with mean between all 3 channels subtracted to highlight the differences between the three channels. Errors in c,d in s.e.m. **e,** Principal components 1-4 (left to right) of the hyperspectral example image shown in Fig. 1j, (cf. Fig. 1b-d). **f,** Achromatic-mean-subtracted mean luminance of the 4 opsin channels from the same scene (like in d). Errors in s.e.m..

**Supplementary Figure 2 | related to Figure 2:**
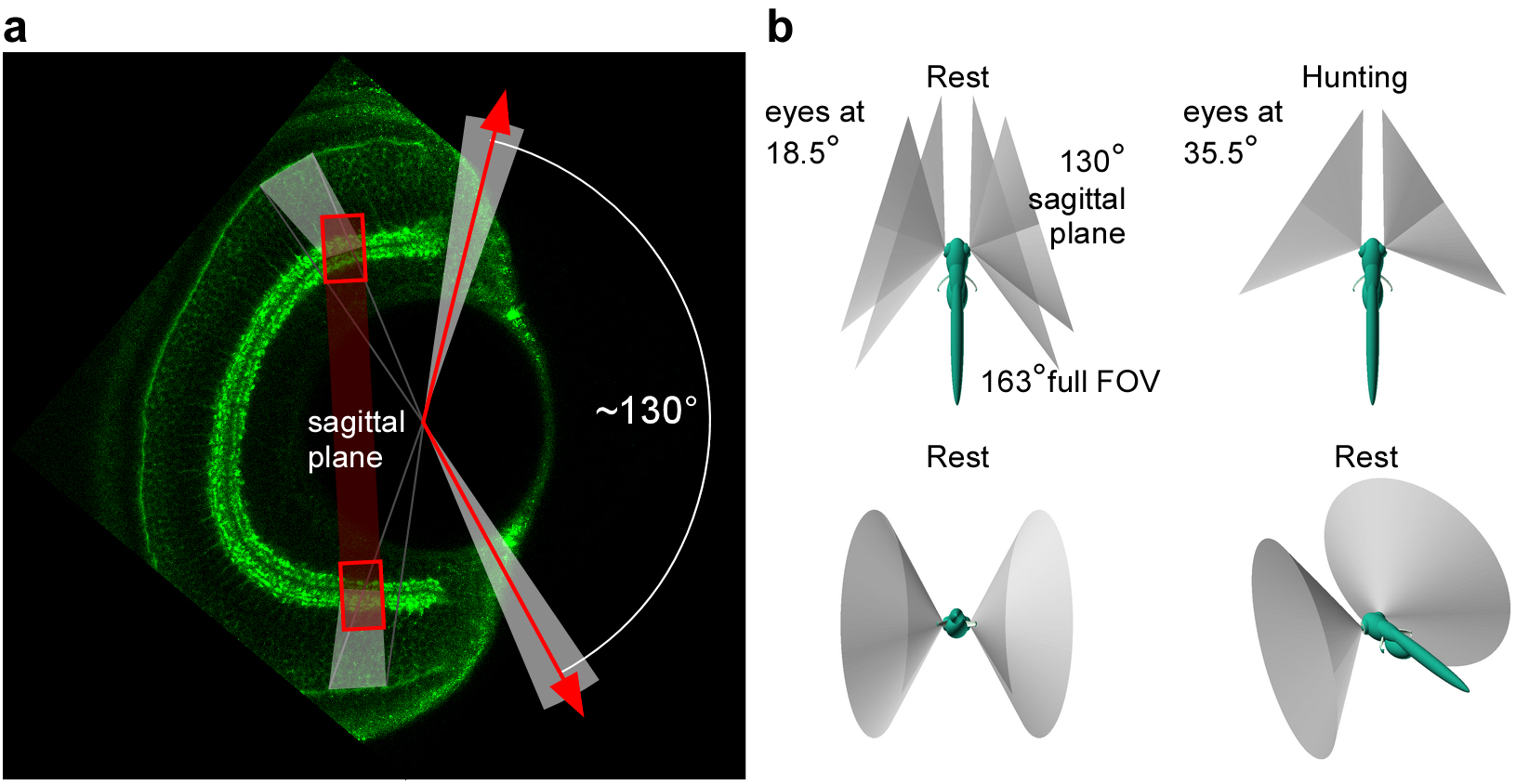
Anisotropic retinal structure. **a,** Transverse plane (looking down onto the fish from the top) immunolabelled 7 *dpf* larval zebrafish eye with bipolar cell terminals in green (like Fig. 2i) highlights the ~130° visual angle surveyed by the sagittal plane used throughout this work (indicated in red). **b,** 3D illustrations of the field of view covered by this sagittal plane during “rest” (eyes tilted forward ~18.5°) and during prey-capture (“hunting”, eyes at 35.5°) as indicated. In the top left panel, the ~163° full field of view of the eye is indicated in addition to the 130° field surveyed in this work.

**Supplementary Figure 3 | related to Figure 3:**
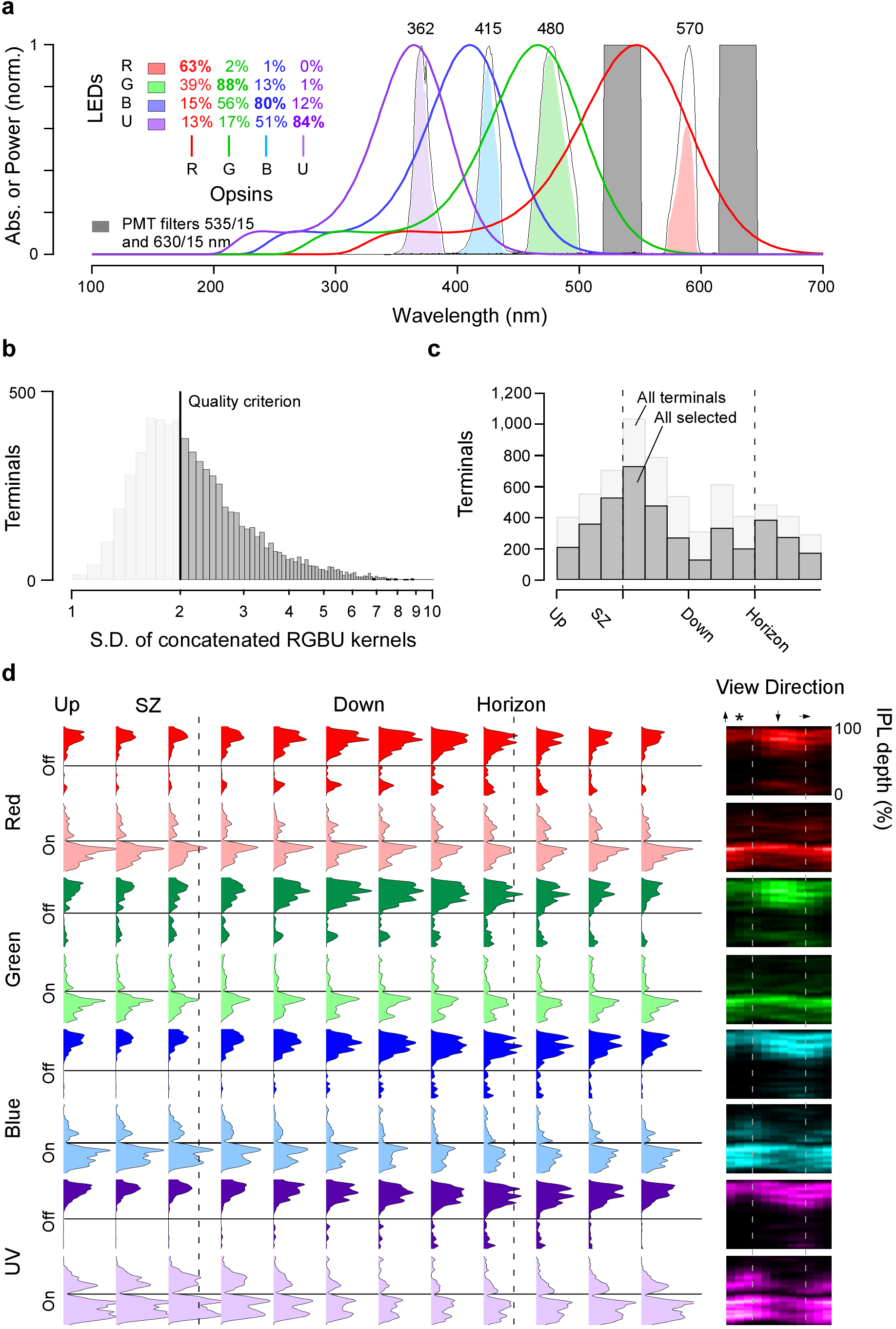
Surveying inner retinal chromatic responses *in vivo*. **a,** Cone action spectra (thin lines) plotted on top of effective LED spectra as measured at the sample plane (black lines) and each LED’s spectral cross-section with respective target cone in solid colours. Grey boxes indicate the positions of the two detector bands. Cross activation efficiencies between LEDs and each cone action spectrum are listed in the inset. **b,** Histogram of quality criterion (Methods) for all recorded terminals with cut-off value of 2 z-scores indicated and **c,** distribution of all (light) and selected (dark) terminals across the eye. **d,** Distribution of R, G, B, U Off- and On-responses across the IPL depth (vertical axis) and eye position (horizontal axis) as indicated (cf. Figs. 3h-k). The insets to the right show the same data plotted as heatmaps.

**Supplementary Figure 4 | related to Figure 6:**
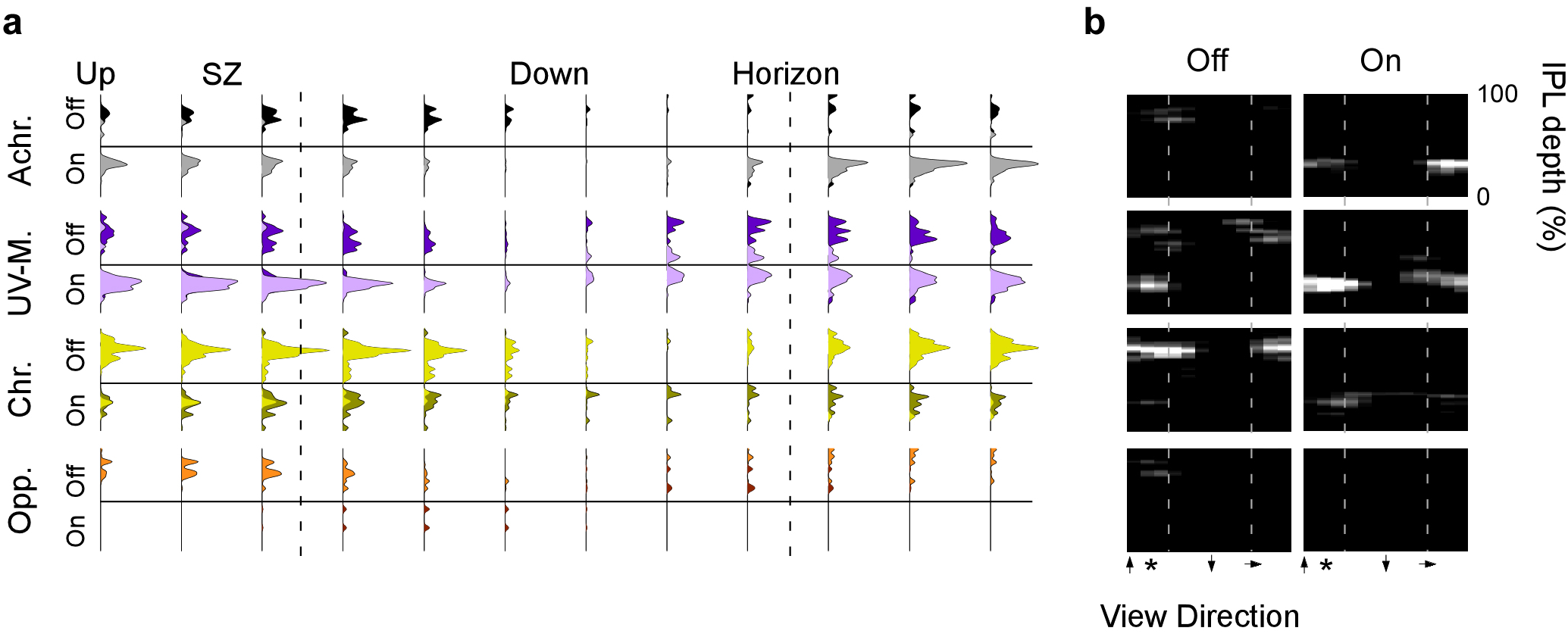
Distribution of genetically defined BC-types. **a,** Distribution of xfz43-positive BC responses split by polarity (Off / On) and allocated major response group plotted against IPL depth (vertical axis) and eye position (horizontal axis). **b**, Same data as (a), plotted as heatmaps.

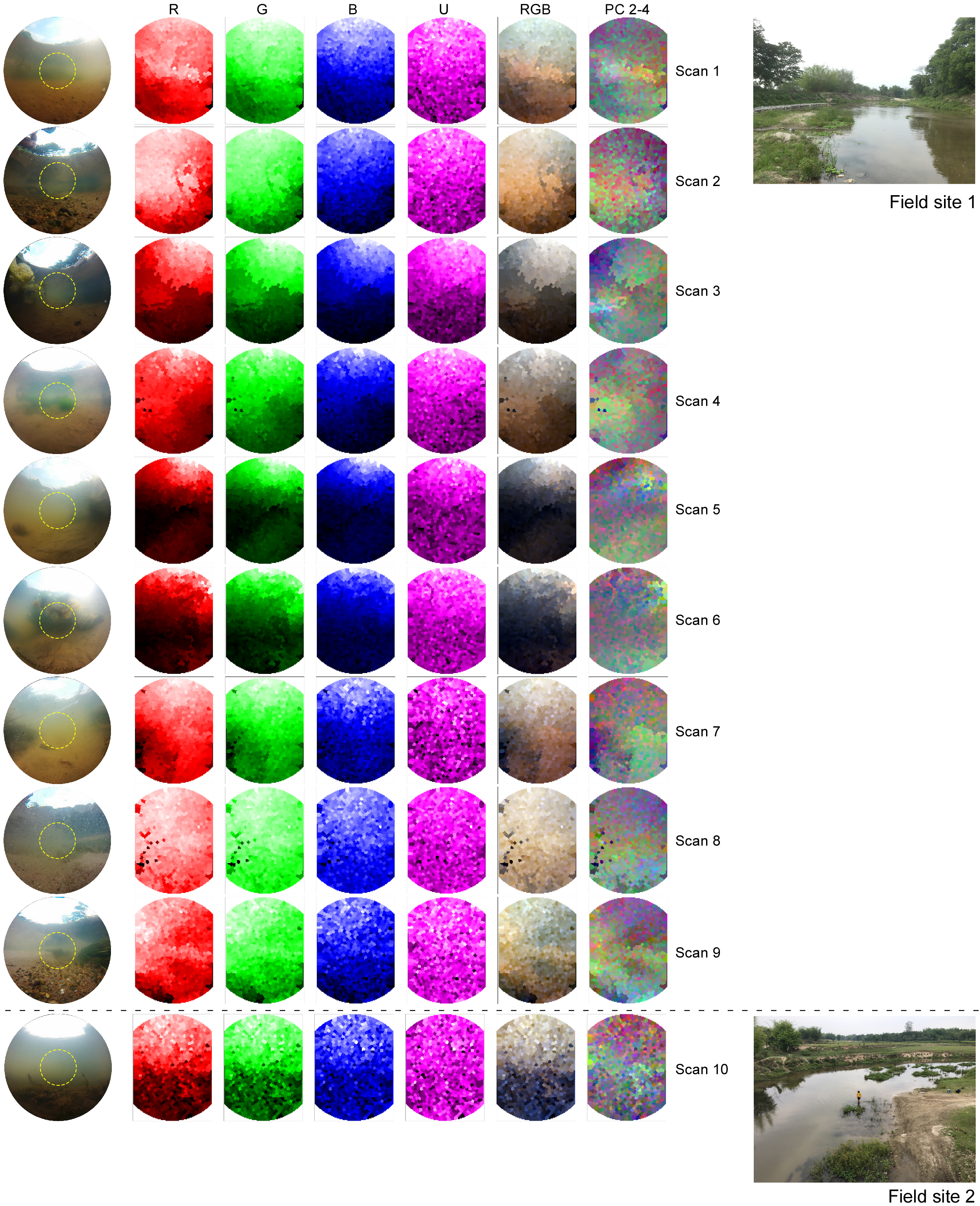
Supplementary Data Sheet1.

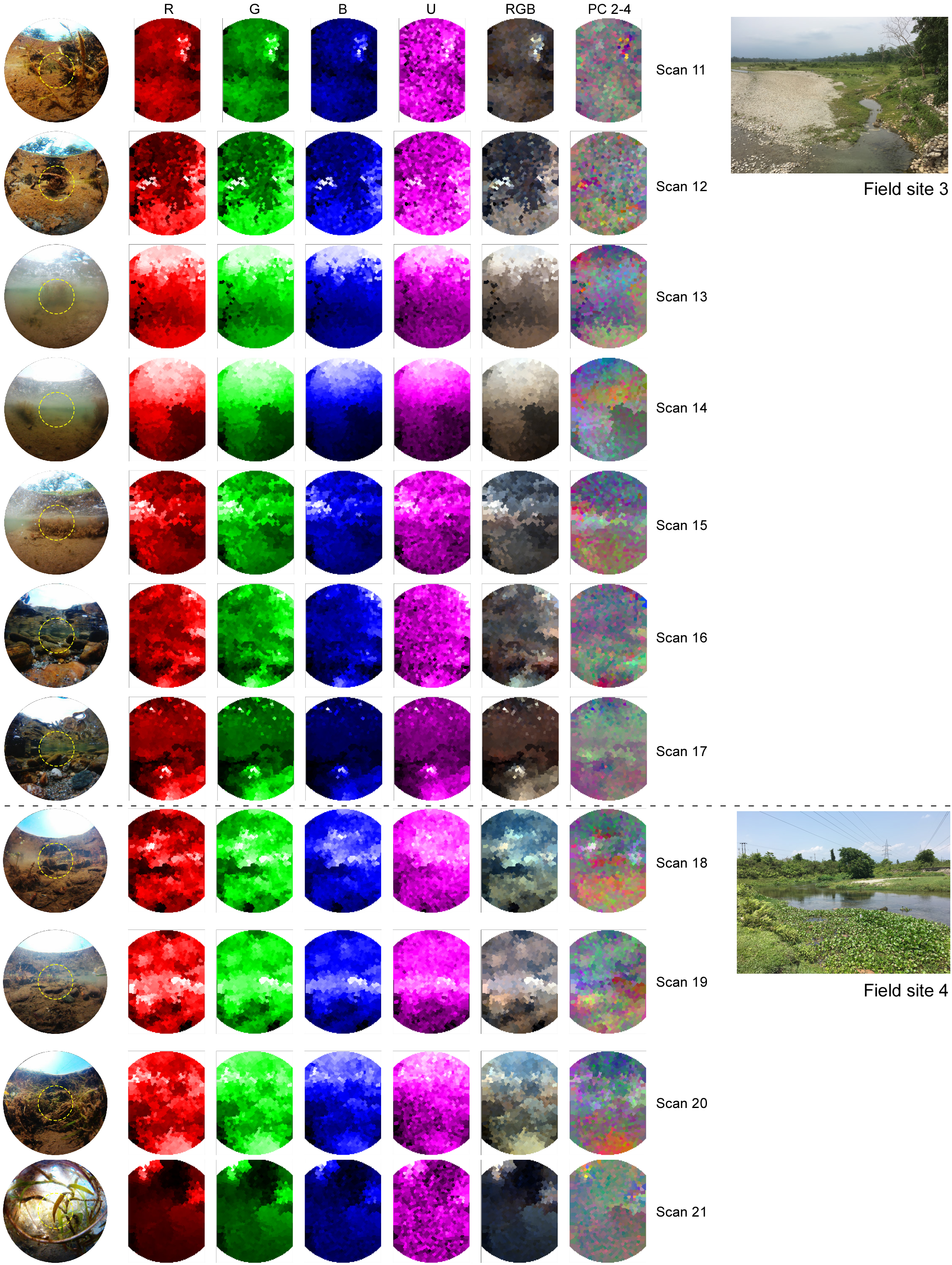
Supplementary Data Sheet2.

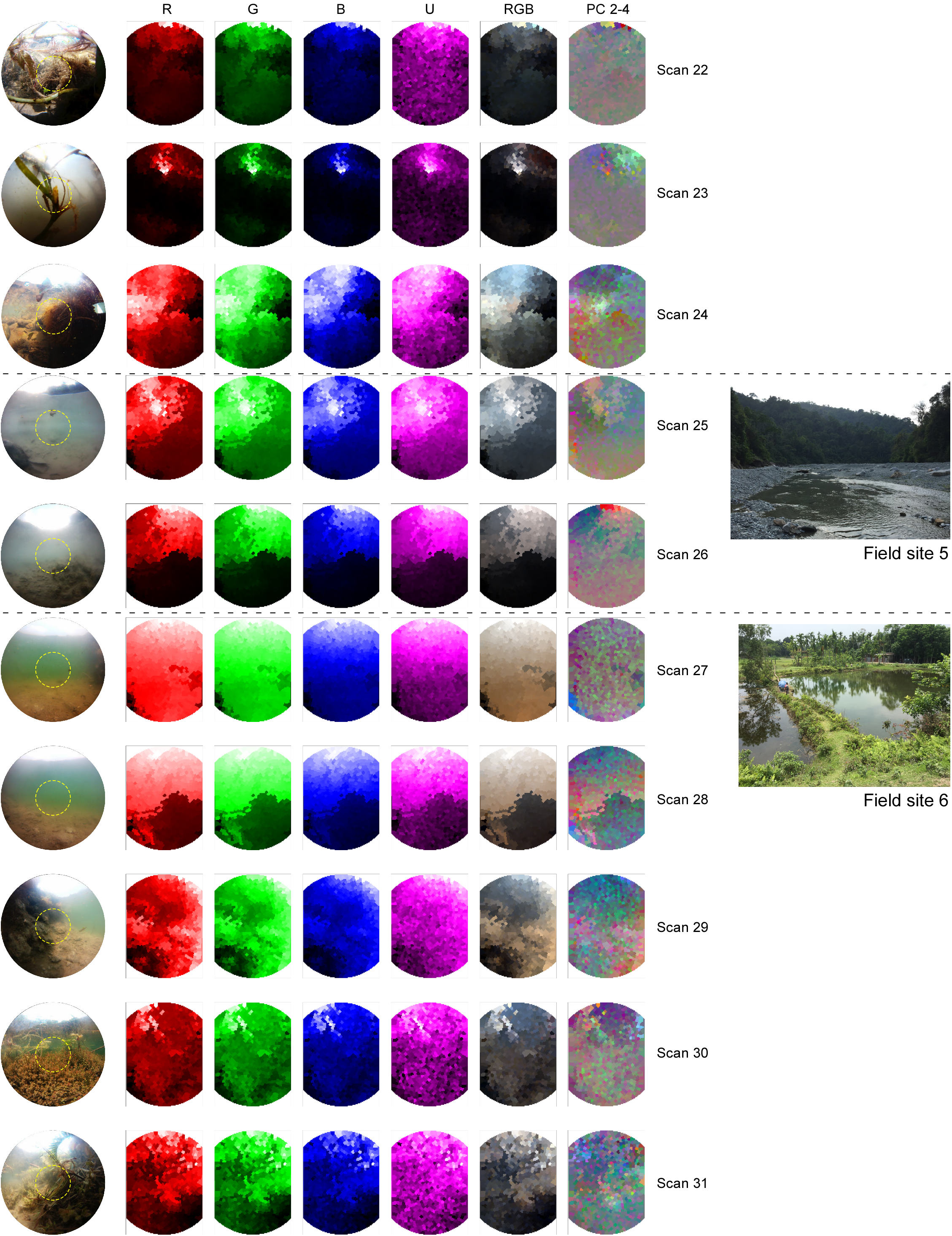
Supplementary Data Sheet3.

## Author contributions

MJY, NEN, TY and TB designed the study, with help from DEN, DO and PB; MJYZ and TY performed 2-photon imaging experiments and immunohistochemistry; TY generated all novel zebrafish lines; NEN and TB built the hyperspectral scanner and performed field work; NEN, MJYZ and TY performed pre-processing with inputs from TB; PB developed the clustering framework; TB analysed the data with help from all authors; TB wrote the manuscript with help from PB and inputs from all authors.

## Acknowledgements

We thank Kripan Sarkar and Fredrik Jutfeld for help with fieldwork, Leon Lagnado for the provision of zebrafish lines and critical feedback and Thomas Euler for critical feedback. The authors would also like to acknowledge support from the FENS-Kavli Network of Excellence. Funding was provided by the European Research Council (ERC-StG “NeuroVisEco” 677687 to TB), Marie Curie Sklodowska Actions individual fellowship (“ColourFish” 748716 to TY), Marie Sklodowska-Curie European Training network “Switchboard” (Switchboard receives funding from the European Union’s Horizon 2020 research and innovation programme under the Marie Sklodowska-Curie grant agreement No. 674901), The Deutsche Forschungsgemeinschaft (DFG: BA 5283/1-1 to TB, BE 5601/1-1 to PB), The Medical Research Council (MC_PC_15071 to TB). The Federal Ministry of Education and Research of Germany through the Bernstein Award for Computational Neuroscience (FKZ 01GQ1601 to PB).

